# *Salmonella* sensitizes macrophages for context-dependent anti-inflammatory responses

**DOI:** 10.64898/2026.01.13.699354

**Authors:** Gloria Alvarado, Keara Lane

**Author notes:** Corresponding author: Keara Lane.

## Abstract

Macrophages’ critical role in the innate immune response depends on their ability to change their activation state in response to environmental signals, generating cells with diverse functions ranging from antimicrobial activity to tissue repair. However, this plasticity in activation state represents a vulnerability that intracellular pathogens have evolved to exploit. *Salmonella* Typhimurium (STm) harnesses macrophage plasticity to promote anti-inflammatory states conducive to bacterial survival and replication. Although this anti-inflammatory state is well-characterized, how it is established remains unclear. Using smFISH, we find that STm is a weak pro-inflammatory trigger and fails to directly induce anti-inflammatory polarization. Instead, we show that STm creates a sensitized cell state with enhanced responsiveness to the cytokine IL-4. While we found that IL-4Rɑ upregulation is a common feature of infected cells, the sensitized state is independent of receptor upregulation. Polarization resulting from this sensitized state is context-dependent, with bacterial load and effector secretion distinctly amplifying expression of the anti-inflammatory genes, *Il4ra* and *Arg1*. Together, our results establish a two-signal model in which pathogen-driven sensitization and environmental IL-4 combine to drive macrophage anti-inflammatory polarization.

## INTRODUCTION

Macrophages are central to host defense, playing key roles in the initiation and resolution of innate immune responses to infection (Murray, 2017). The functional diversity of macrophages relies on their ability to adjust their gene expression profile and activation state in response to inflammatory signals in the environment (Stout *et al*., 2005; Lawrence and Natoli, 2011; Smith *et al*., 2016; Liu *et al*., 2020). For example, lipopolysaccharide (LPS) and interferon-γ (IFN-γ) drive M1 polarization, which is characterized by pro-inflammatory cytokine production and antimicrobial responses (Mills *et al*., 2000; Eckmann and Kagnoff, 2001). IL-4 and IL-10 induce M2 polarization, which is involved in the resolution of infection and tissue repair (Martinez *et al*., 2009; Bosurgi *et al*., 2017). M1 and M2 responses define opposing ends of macrophage polarization. However, during infection these responses are more diverse, spanning the continuum of polarization states (Smith *et al*., 2016; Palma *et al*., 2018). While the ability to adjust activation state allows macrophages to generate tailored responses to signals in their local environment, it also represents a vulnerability that pathogens have evolved to exploit (Shirey *et al*., 2008; Jaslow *et al*., 2018). Understanding how infections alter macrophage responses to polarizing signals is therefore critical to determine how inflammatory responses are generated during infection.

*Salmonella enterica* serovar Typhimurium (STm) is an intracellular bacterial pathogen that manipulates macrophage gene expression to survive (Monack *et al*., 2004; Helaine *et al*., 2014; Stapels *et al*., 2018). Although STm induces pro-inflammatory responses via TLR4 activation, STm was found to predominantly reside in anti-inflammatory macrophages during chronic infections in mice, identifying this macrophage activation state as a permissive environment for STm (Eisele *et al*., 2013). However, understanding how macrophages transition to this anti-inflammatory state during infection has been stymied by the heterogeneity of bacterial infections. Even in cell culture models of infection, extensive variability in infection outcome is observed (Helaine *et al*., 2010; Avraham *et al*., 2015). Only a subset of exposed macrophages become infected, and even among infected cells, bacterial replication is variable. The advent of single-cell RNA-seq (scRNA-seq) enabled the characterization of these heterogeneous macrophage subpopulations during STm infection and revealed marked differences in gene expression patterns and activation states (Avraham *et al*., 2015; Saliba *et al*., 2016; Stapels *et al*., 2018; Heyman *et al*., 2023). While uninfected bystanders (cells exposed to but not infected by STm) or those containing non-replicating bacteria had pro-inflammatory transcriptional profiles, macrophages containing replicating STm typically had anti-inflammatory transcriptional profiles. These studies established that heterogeneity in macrophage polarization is associated with diverse bacterial fates and bolstered the link between anti-inflammatory polarization and STm survival and replication.

Single-cell analyses of macrophages during STm infection have revealed how this pathogen affects inflammatory gene expression programs. First, heterogeneity in STm LPS modifications has been shown to correlate with type 1 IFN responses. Strong IFN induction only occurs in a subset of infected cells (Avraham *et al*., 2015). Second, intracellular STm manipulates macrophage signaling and activation state by expressing the *Salmonella* Pathogenicity Island 2 (SPI-2) virulence regulon, which encodes a type 3 secretion system (T3SS) that secretes effectors into the host cytoplasm (LaRock *et al*., 2015). These effectors inhibit pro-inflammatory signaling proteins, including the transcription factor NF-κB (Rolhion *et al*., 2016; Jennings *et al*., 2017). Anti-inflammatory signaling is also a target of STm effectors, with SteE-mediated activation of STAT3 leading to the transcription of anti-inflammatory genes, such as *Il4ra* (LaPorte *et al*., 2008; Jaslow *et al*., 2018; Stapels *et al*., 2018; Gibbs *et al*., 2020; Panagi *et al*., 2020). Anti-inflammatory gene expression responses have also been found to be positively correlated with bacterial load in macrophages (Heyman *et al*., 2023). Together, these results demonstrate that STm can drive changes in macrophage gene expression and activation states.

During infection, macrophage polarization responses occur in dynamic cytokine environments. Infected macrophages release chemokines to recruit additional immune cells to the site of infection (Mantovani *et al*., 2004). Some of these cells are antimicrobial (Eckmann and Kagnoff, 2001), while others promote the resolution of inflammation by releasing anti-inflammatory cytokines (Bosurgi *et al*., 2017). How macrophages integrate complex mixtures of polarization signals to generate a robust inflammatory response is incompletely understood. Several studies support either mutual inhibition between M1 and M2 polarization states (Modolell *et al*., 1995; Ohmori and Hamilton, 1997; Smallie *et al*., 2010; Piccolo *et al*., 2017; Zhang and Jagannath, 2025) or mixed responses upon simultaneous exposure to both types of polarization signals (Smith *et al*., 2016; Muñoz-Rojas *et al*., 2021). The sequence of polarization signal exposure may also affect activation state, with prior exposure to one signal increasing responsiveness upon subsequent exposure to the opposing polarization signal (D’Andrea *et al*., 1995; Stout *et al*., 2005; Czimmerer *et al*., 2022). For example, macrophages pre-stimulated with LPS and IFN-γ were found to have enhanced anti-inflammatory gene expression when subsequently treated with IL-4, compared to cells treated with IL-4 alone (O’Brien and Spiller, 2022). Although multiple distinct mechanisms can be responsible for M2 polarization, how the transition to an M2 polarized state is established during STm infection remains unclear. For instance, it is unknown whether pathogen-derived signals are sufficient to directly induce anti-inflammatory polarization, or whether infection sensitizes macrophages to M2 polarizing signals that then trigger polarization.

Here, we use single-molecule fluorescent in situ hybridization (smFISH) to investigate how macrophage polarization states are established during STm infection. Using conditionally immortalized macrophages (CIMs) as our model system (Roberts *et al*., 2019), we find that STm is a weak trigger of pro-inflammatory responses and fails to directly induce anti-inflammatory responses. Instead, STm creates a sensitized macrophage state characterized by enhanced responsiveness to IL-4 that is largely independent of *Il4ra* upregulation. Bacterial load and effector secretion modulate the degree of sensitization, amplifying the gene expression response to IL-4. Together, our results establish a two-signal model for anti-inflammatory polarization during STm infection of CIMs in which pathogen-driven sensitization and environmental IL-4 combine to drive a spectrum of macrophage anti-inflammatory states.

## RESULTS

### *Salmonella* induces heterogeneous and divergent *Tnf* and *Nos2* gene expression responses

To investigate how anti-inflammatory polarization of macrophages is established during STm infection, we used conditionally immortalized macrophages (CIMs) as our model system (Roberts *et al*., 2019). CIMs are derived from myeloid progenitors immortalized through conditional expression of Hoxb8; removal of Hoxb8 and addition of M-CSF enables macrophage differentiation. This approach generates a renewable progenitor stock that serves as a continuous source of macrophages. These cells and similar derivatives have been found to mimic key features of primary bone marrow-derived macrophages (BMDMs), including similar gene expression profiles, cytokine production, supporting the growth of intracellular pathogens such as *Mycobacterium tuberculosis (Mtb)*, and limited lifespan post-differentiation (Knoepfler *et al*., 2001; Wang *et al*., 2006; Roberts *et al*., 2019; Luecke *et al*., 2024). CIMs have several advantages for studying anti-inflammatory polarization compared to transformed macrophage-like cell lines, such as RAW264.7 cells. Anti-inflammatory polarization requires oxidative phosphorylation, but transformed cell lines rely on aerobic glycolysis, making them poorly suited for such studies (Cairns *et al*., 2011; He *et al*., 2021; Zhang and Jagannath, 2025). In contrast, CIMs are expected to more closely mimic the metabolic profile of BMDMs due to their finite lifespan and non-transformed state. Therefore, CIMs are a valuable system to dissect anti-inflammatory responses to STm, marrying the experimental advantages of immortalized cells with physiologically relevant responses.

To quantify pro-inflammatory responses to STm, we infected CIMs with fluorescent STm (mCerulean3) and also treated cells with M1-polarizing ligands (LPS and IFN-γ) as a benchmark for comparison (Figure 1A). Macrophage responses are highly heterogeneous; thus, we used single-molecule fluorescent *in situ* hybridization (smFISH) rather than scRNA-Seq to measure gene expression in individual macrophages, as it is ideally suited to identify cells present in low proportions, and avoids issues related to poor detection or transcriptome coverage (Torre *et al*., 2018). Because smFISH is a lower-throughput technique, we measured gene expression of two canonical pro-inflammatory markers, *Tnf* and *Nos2*, at 4 hours and 24 hours post-infection (Figures 1B, S1.1). To distinguish responses driven by extracellular exposure from those driven by uptake and intracellular infection, we segmented fluorescent bacteria in the images and classified CIMs as infected or uninfected based on the presence or absence of intracellular bacterial objects, respectively (see Methods). M1 ligands induced robust *Tnf* expression at 4 hours that was maintained through 24 hours, with nearly all CIMs responding. STm induced a similar response, but with weaker overall *Tnf* expression in both infected and uninfected cells. This indicates that extracellular bacterial exposure, rather than intracellular infection, is responsible for the *Tnf* response. In contrast to *Tnf,* the *Nos2* expression response differed between ligands and STm. M1 ligands induced robust *Nos2* expression at 4 hours in the majority of cells, with per cell expression levels increasing through 24 hours. In contrast, STm induced *Nos2* expression in a much smaller fraction of cells, and the time-dependent increase observed with M1 ligands was predominantly observed in infected rather than uninfected cells.

**Figure 1.**
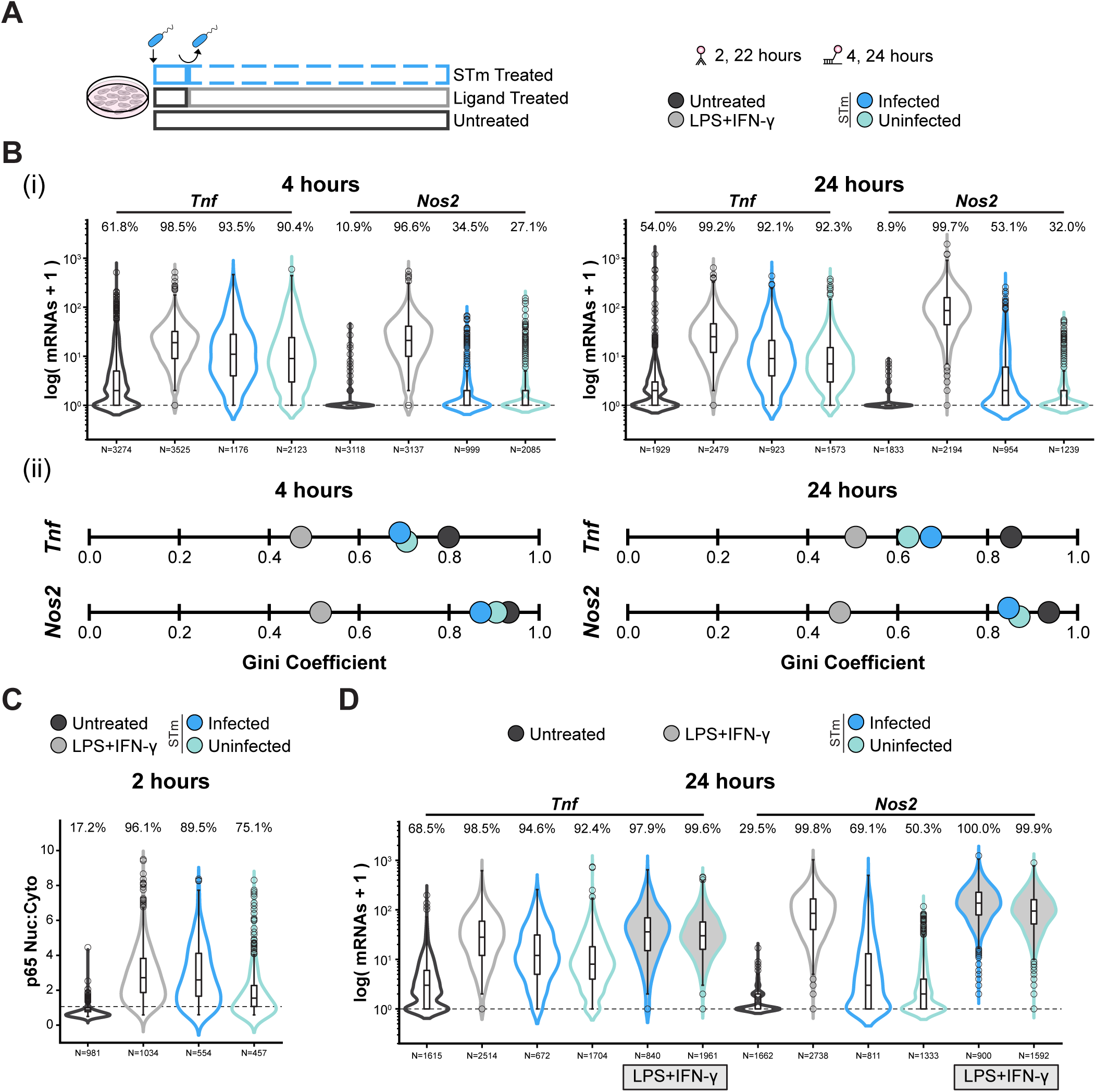
STm induces divergent *Tnf* and *Nos2* gene expression in CIMs. (A) Schematic of the experimental setup. In (B) and (C), CIMs were left untreated, treated with M1 ligands (100 ng/mL LPS and 5 ng/mL IFN-γ), or exposed to STm (MOI 10). Samples were fixed at the indicated timepoints, processed for smFISH or immunofluorescence (IF), and imaged. CIMs were classified as infected or uninfected based on the presence or absence of segmented bacterial objects, respectively (see Methods). (B) (i) smFISH shows STm induces *Tnf* but limited *Nos2* expression. Violin plots show transcript levels for *Tnf* and *Nos2* after stimulation for 4 or 24 hours with the indicated treatment. Data are presented as log(mRNA +1). Dashed line indicates expressing CIMs and the percentage of CIMs containing transcripts are above the violins. (ii) Gini coefficients quantifying expression heterogeneity for data in (i). Gini coefficient of 0 indicates perfect equality and 1 indicates perfect inequality. (C) STm induces p65 signaling. p65 localization at 2 hours measured by IF. Signal is displayed as the p65 nuclear:cytoplasmic ratio of median intensity values (p65 Nuc:Cyto). Percentages above violins indicate the percentage of CIMs with p65 signal above the baseline (dashed line, defined as 1 SD above the mean of untreated CIMs, see Figure S1.3 and Methods). (D) STm exposed CIMs respond to M1 ligands. *Tnf* and *Nos2* expression measured at 24 hours using smFISH. Conditions and plotting details are as in (B) (i) with the addition of STm-exposed CIMs treated with M1 ligands (filled in violins). Data in (B), (C), and (D) are pooled from three biological replicates, N=total number of CIMs. The width of violins represents the proportion of CIMs within the respective condition. Gini coefficients and the results of statistical analyses on smFISH data are reported in Table S1.

Heterogeneity is a hallmark of macrophage transcriptional responses, enabling distinct functional outcomes in a population of cells exposed to a uniform stimulus (Avraham *et al*., 2015; Xue *et al*., 2015; Lane *et al*., 2017; Bagnall *et al*., 2018; Muñoz-Rojas *et al*., 2021). To quantify heterogeneity in gene expression, we calculated the Gini coefficient (Figure 1B, Table S1), which is a measure of inequality that ranges from 0 (all cells express the same number of transcripts) to 1 (all transcripts originate from a single cell) (Shaffer *et al*., 2017). For both *Tnf* and *Nos2*, Gini coefficients were higher in STm-exposed cells (*Tnf*: 0.62-0.71; *Nos2*: 0.85-0.91) than in ligand-treated cells (*Tnf*: 0.47-0.51; *Nos2*: 0.47-0.51), indicating that STm induces more heterogeneous gene expression responses than purified ligands. Together, these results demonstrate that the pro-inflammatory response of CIMs to STm is distinct from responses to purified ligands. While *Tnf* expression is robustly induced by extracellular bacterial exposure, *Nos2* expression is minimally induced and restricted mainly to infected cells. This reveals that STm induces a divergent pro-inflammatory response, where robust *Tnf* expression is accompanied by muted *Nos2* induction in a select fraction of infected cells.

### STm weakly induces *Nos2* despite robust NF-κB activation

The muted *Nos2* response to STm (Figure 1B) could arise through several mechanisms. First, CIMs might express low levels of TLR4, making IFN-γ rather than LPS primarily responsible for *Nos2* expression in response to M1 ligands. Second, STm might weakly activate TLR4 and its downstream signaling pathways, including NF-κB, the major transcription factor for *Nos2* (Xie *et al*., 1994). Third, STm might actively suppress TLR4/NF-κB signaling through secreted effector proteins. To determine whether CIMs express *Nos2* in response to a purified TLR4 ligand, we treated CIMs with LPS alone, LPS and IFN-γ, or STm and used smFISH to quantify *Nos2* transcripts at 4 and 24 hours post-infection (Figure S1.2). CIMs expressed *Nos2* in response to LPS alone, although at reduced levels compared to combined LPS and IFN-γ treatment. This demonstrates that TLR4 signaling is intact and is sufficient to induce *Nos2* expression. To test whether STm weakly activates TLR4 signaling, we used immunofluorescence to quantify nuclear NF-κB (p65 subunit) at 2 and 22 hours post-infection. Timepoints were chosen to correspond with smFISH experiments and account for signaling preceding transcription. At 2 hours, the majority of cells across all treatments displayed nuclear NF-κB (Figures 1C, S1.3A). Unlike *Tnf*, which is a primary response gene and thus rapidly activated, *Nos2* is a secondary response gene, thus, its expression is slower because it requires new protein synthesis (Ramirez-Carrozzi *et al*., 2006; Tong *et al*., 2016). However, NF-κB nuclear localization was maintained at 22 hours across all treatments (Figure S1.3B). This indicates that the muted *Nos2* response to STm is not due to impaired NF-κB activation. Finally, to determine whether STm actively suppresses TLR4 signaling and *Nos2* expression, we treated STm-exposed cells with M1 ligands. If STm secrete effectors that inhibit *Nos2* expression, the addition of M1 ligands should not rescue *Nos2* expression in infected cells. Instead, we observed that cells exposed to STm and treated with M1 ligands induced *Nos2* to levels comparable to ligands alone, regardless of infection status (Figures 1D, S1.4). Similar responses were observed for *Tnf*. These results indicate that STm does not suppress *Nos2* induction and that cells remain responsive to M1 ligands. Taken together, these results demonstrate that STm is a weak pro-inflammatory trigger for *Nos2* expression and that the basis for the muted *Nos2* response lies downstream of NF-κB activation.

### The muted pro-inflammatory response to STm is not due to anti-inflammatory polarization

Macrophage pro- and anti-inflammatory transcriptional programs are often considered mutually exclusive, with studies supporting mutual cross-inhibition between the two states (Modolell *et al*., 1995; Smallie *et al*., 2010; Piccolo *et al*., 2017; Zhang and Jagannath, 2025). Intracellular STm have recently been shown to promote an anti-inflammatory phenotype through effector secretion (Jaslow *et al*., 2018; Stapels *et al*., 2018; Gibbs *et al*., 2020; Panagi *et al*., 2020). Therefore, we hypothesized that the weak pro-inflammatory response induced by STm might result from the activation of an anti-inflammatory program that inhibits pro-inflammatory gene expression. To test this hypothesis, we quantified anti-inflammatory responses to STm. We infected CIMs with fluorescent STm and also treated CIMs with M2-polarizing ligands (IL-4 and IL-10) as a benchmark for anti-inflammatory gene expression. We used smFISH to measure gene expression of two canonical anti-inflammatory marker genes, *Mrc1* and *Arg1*, at 4 hours and 24 hours post-infection (Figures 2A, S2.1). STm failed to induce expression of either anti-inflammatory marker gene at 4 or 24 hours. As expected, M2 ligands induced robust *Mrc1* expression at 4 hours, which further increased by 24 hours. Unexpectedly, M2-ligand stimulation failed to induce robust *Arg1* expression. Even among the subset of cells (51%) with expression at 24 hours, *Arg1* transcript levels remained low (mean = 20 transcripts). To quantify response heterogeneity, we calculated Gini coefficients for each condition (Figure 2A, Table S1). Gini coefficients were high (0.82-0.90) for both genes in all conditions, with the exception of *Mrc1* in M2 ligand-treated cells (∼0.46), indicating that anti-inflammatory gene expression in CIMs is highly heterogeneous.

**Figure 2.**
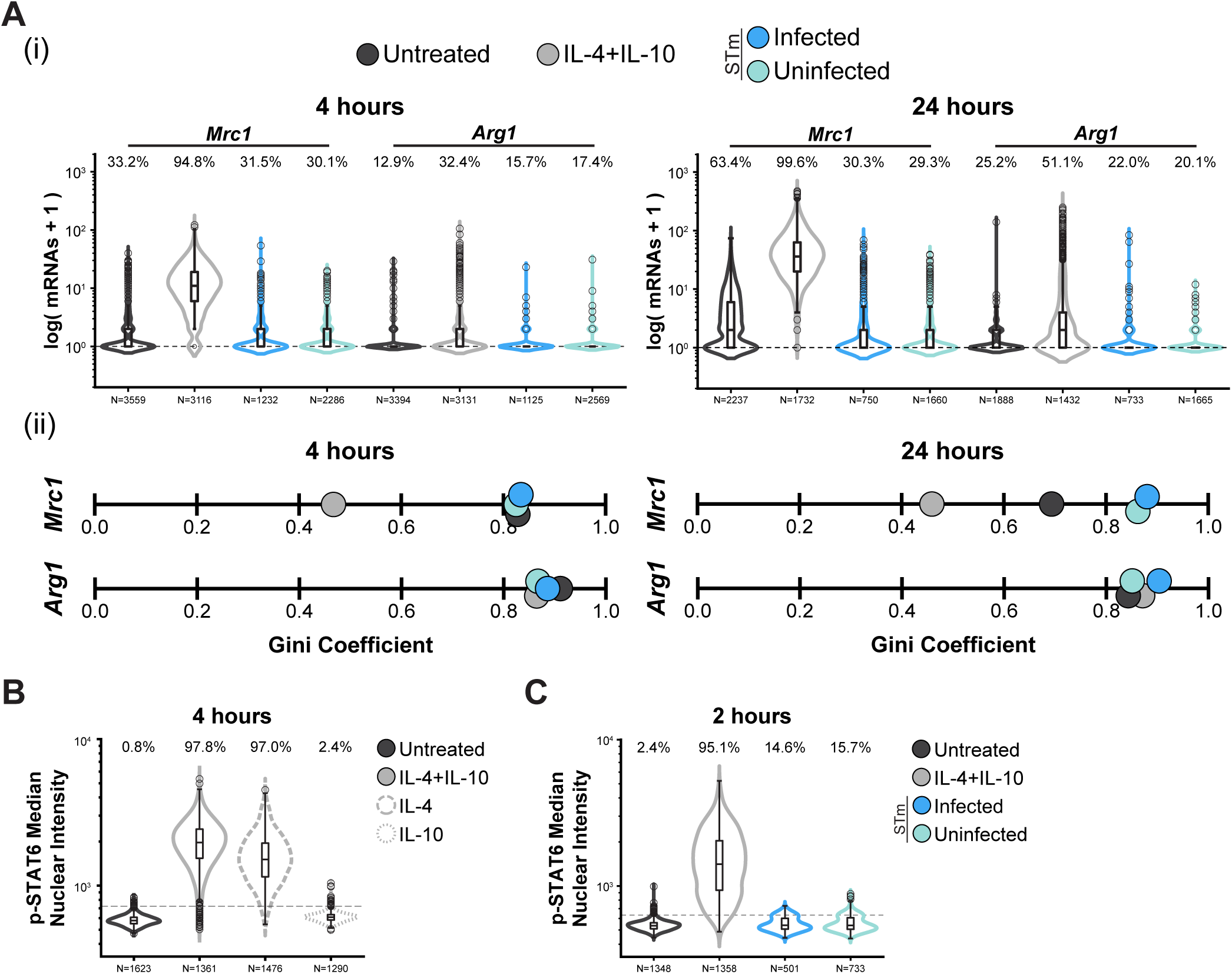
STm fails to induce robust anti-inflammatory gene expression and signaling in CIMs. (A) (i) STm does not induce expression of anti-inflammatory marker genes (*Mrc1* and *Arg1*). CIMs were left untreated, treated with M2 ligands (25 ng/mL IL-4 and 25 ng/mL IL-10), or exposed to STm (MOI 10). Samples were fixed at 4 and 24 hours, processed for smFISH, and imaged. Violin plots show transcript levels for *Mrc1* and *Arg1*; data are presented as log(mRNA +1). Dashed line indicates expressing CIMs and the percentage of CIMs containing transcripts are above the violins. (ii) Gini coefficients quantifying expression heterogeneity for data in (i). (B) IL-4 but not IL-10 drives STAT6 phosphorylation in CIMs. CIMs were left untreated or treated with IL-4 (25 ng/mL), IL-10 (25 ng/mL), or both ligands. Samples were fixed at 4 hours, stained for p-STAT6, and imaged. Data from a representative experiment (N=3 total) is shown. Violin plots show median nuclear p-STAT6 intensity. Percentages above violins indicate the percentage of CIMs with p-STAT6 above the basal signal threshold (dashed line). (C) STm does not induce STAT6 phosphorylation. CIMs were left untreated, treated with M2 ligands, or exposed to STm. Samples were fixed at 2 hours, stained for p-STAT6, and imaged. Data is plotted as in (B). The basal signal threshold for (B) and (C) is defined as described in Figure S2.2B and Methods. Data in (A) and (C) are pooled from three biological replicates, N=total number of CIMs. CIMs were classified as infected or uninfected as in Figure 1 (see Methods). The width of violins represents the proportion of CIMs within the respective condition. Gini coefficients and the results of statistical analyses on smFISH data are reported in Table S1.

Although induction of the anti-inflammatory gene *Arg1* in response to M2 ligands is known to be variable, with many non-expressing cells (Muñoz-Rojas *et al*., 2021), we wanted to confirm that the lack of a robust *Arg1* response in CIMs could not be explained by low or heterogeneous expression of the receptors for the M2 ligands. Receptor activation, primarily via IL-4 (LaPorte *et al*., 2008), induces STAT6 phosphorylation, which subsequently translocates to the nucleus and initiates transcription of *Mrc1* and *Arg1*. Thus, to determine whether CIMs can activate signaling in response to M2 ligands, we treated cells with the ligands and measured nuclear phospho-STAT6 (p-STAT6) at 4 hours using immunofluorescence (Figures 2B, S2.2A). The majority of cells displayed nuclear p-STAT6, and as expected, IL-4 is the primary driver of STAT6 phosphorylation. This demonstrates that CIMs are uniformly competent to respond to IL-4 and that the weak *Arg1* response to M2 ligands is not due to impaired upstream signaling. In contrast, STm-exposure weakly activates STAT6 (Figures 2C, S2.2B-C), and this response occurs in only a small subset of cells (∼15%) at both early (2 hours) and late (22 hours) stages of infection, with similar fractions of responders observed in both infected and uninfected cells. This indicates that STm does not robustly trigger STAT6 signaling, regardless of whether bacteria are phagocytosed. Thus, while both M2 ligands and STm do not robustly induce *Arg1* in CIMs, the underlying mechanisms for the failed response are distinct. Taken together, these data demonstrate that STm does not induce a canonical anti-inflammatory transcriptional response in CIMs. The muted pro-inflammatory response to STm, therefore, cannot be the result of mutual inhibition between polarization states.

### *Il4ra* is highly upregulated in a subset of *Salmonella*-infected macrophages

The low-level activation of STAT6 (∼15% cells with low signal intensity) (Figures 2C, S2.2B-C) suggested that STm initiates a weak anti-inflammatory response in a subset of cells. However, the absence of *Mrc1* and *Arg1* expression indicates that this response is insufficient to induce expression of canonical anti-inflammatory marker genes. To further investigate this weak anti-inflammatory response, we examined *IL4ra* gene expression. As the receptor at the apex of the IL-4 signaling pathway, *Il4ra* expression acts as an early marker of the anti-inflammatory response (Kotanides and Reich, 1996), and thus could reveal the extent to which a partial anti-inflammatory response is initiated. Furthermore, *Il4ra* has been shown to be upregulated in anti-inflammatory macrophages containing STm (Saliba *et al*., 2016; Stapels *et al*., 2018; Heyman *et al*., 2023). We measured *Il4ra* expression using smFISH (Figures 3A, S3.1A). Approximately 50% of untreated cells had basal *Il4ra* expression. Both M2 ligand treatment and STm exposure increased *Il4ra* expression levels in the majority of cells (76.7-90%) with moderate variability in expression (>0.6) across all conditions as measured by the Gini coefficient (Figure S3.1B, Table S1). However, STm infection resulted in the largest fraction of cells (90%) expressing *Il4ra* and the highest mean expression level per cell at 24 hours (Table S1).

**Figure 3.**
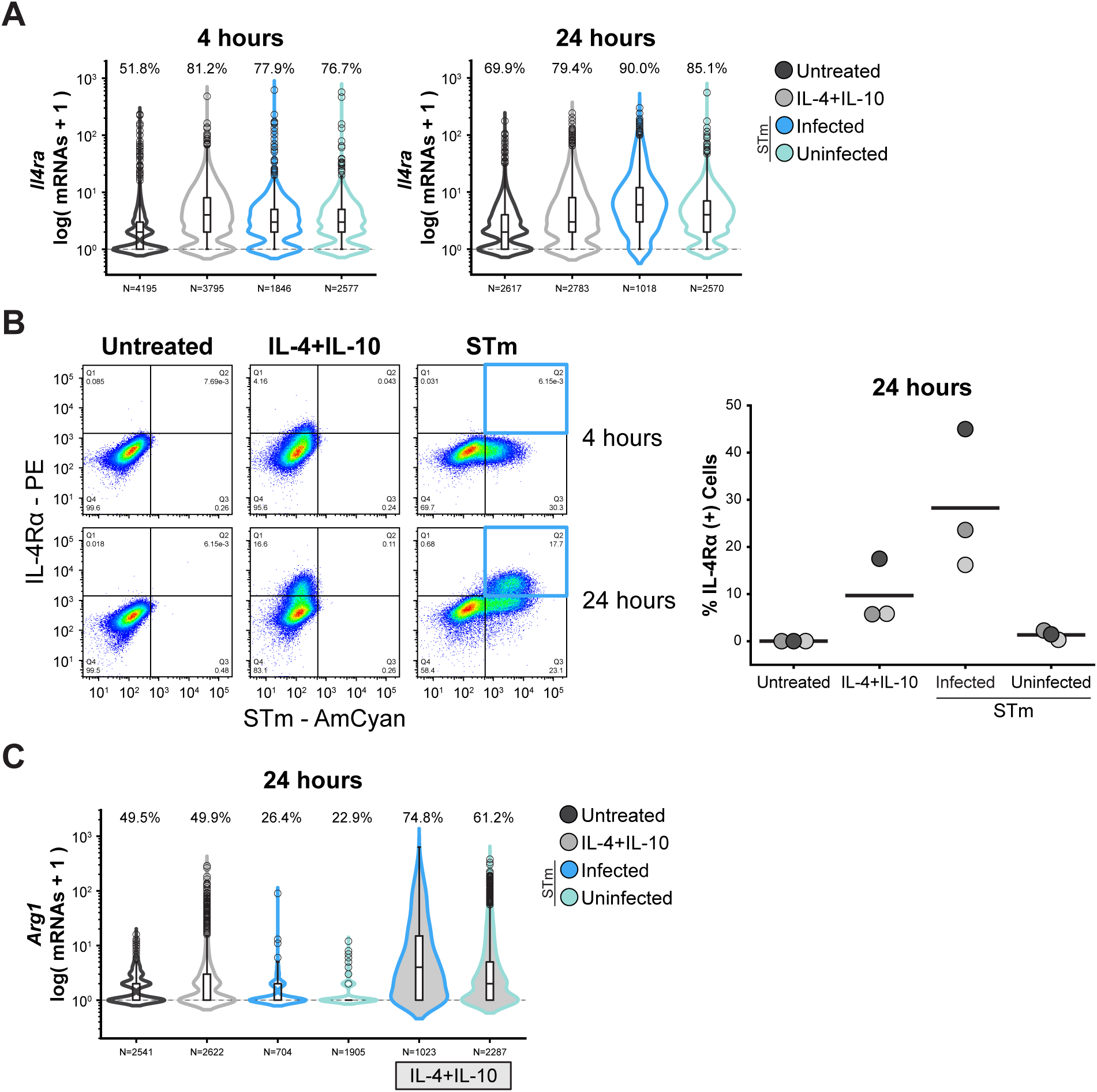
STm sensitizes CIMs to anti-inflammatory ligands. (A) *Il4ra* gene expression increases during STm infection. CIMs were left untreated, treated with M2 ligands (25 ng/mL IL-4 and 25 ng/mL IL-10), or exposed to STm (MOI 10). Samples were fixed at 4 and 24 hours, processed for smFISH, and imaged. Violin plots show transcript levels for *Il4ra*. Data are presented as log(mRNA +1). Dashed line indicates expressing CIMs and the percentage of CIMs containing transcripts are above the violins. (B) A subpopulation of STm-infected CIMs have increased IL-4Rɑ surface protein levels. CIMs were treated as in (A). IL-4Rɑ surface expression (IL-4Rɑ-PE) and STm (STm-AmCyan) were measured by flow cytometry. Representative dot plots from a single experiment (N=3 total) display 65,000 CIMs per sample, percentages included for each quadrant. IL-4Rɑ and STm gates were set using untreated CIMs stained for IL-4Rɑ surface expression. The fraction of IL-4Rɑ+ CIMs for each condition was quantified; biological replicates are shown in shades of gray (dark gray corresponds to data shown in the dot plot), black bars indicate the mean. For STm infected and uninfected CIMs, the fraction of IL-4Rɑ+ CIMs was quantified for each subpopulation separately. As noted in the text, our flow cytometry measurements capture only CIMs with IL-4Rɑ expression above a basal expression level due to limitations in detection. (C) M2 ligand treatment during infection induces *Arg1* expression. CIMs were left untreated, treated with M2 ligands, exposed to STm (empty violins), or exposed to STm followed by treatment with M2 ligands (filled violins)*. Arg1* expression was measured at 24 hours using smFISH. Plotting details are as in (A). Data in (A) and (C) are pooled from three biological replicates. CIMs were classified as infected or uninfected as in Figure 1 (see Methods), N=total number of CIMs. The width of violins represents the proportion of CIMs within the respective condition. Results of statistical analyses on smFISH data are reported in Table S1.

IL-4Rɑ upregulation has functional consequences for macrophage inflammatory state, enabling responses to anti-inflammatory ligands in the environment. We asked whether increased *Il4ra* expression translated to increased receptors on the cell surface. We measured surface IL-4Rɑ protein by flow cytometry in untreated cells, M2 ligand-treated cells, and STm-exposed cells. Given the reduced sensitivity of flow cytometry compared to microscopy, we used an STm strain expressing mCerulean3 from a high-copy plasmid to facilitate identification of infected cells. Despite our earlier observation that IL-4, the ligand for IL-4Rɑ, induces p-STAT6 in the majority of cells (97.8%) (Figure 2B), the IL-4Rɑ-PE signal from untreated cells and the PE signal from isotype controls were indistinguishable by flow cytometry (Figure S3.2A). This indicates that basal IL-4Rɑ levels, while sufficient to activate downstream signaling in response to IL-4, fall below the limit of detection of flow cytometry. Therefore, our flow cytometry measurements capture only cells with IL-4Rɑ expression above this baseline detection level. We observed a consistent increase in surface IL-4Rɑ in a fraction of CIMs treated with M2 ligands or exposed to STm compared to untreated cells (Figures 3B, S3.2B). In STm-exposed cells, increased surface IL-4Rɑ was predominantly associated with infected rather than uninfected cells at 24 hours (Figures 3B, S3.2B). This indicates that intracellular infection, rather than extracellular exposure, drives upregulation of IL-4Rɑ on the cell surface, despite increased *Il4ra* gene expression in both groups of cells. Together, our results demonstrate that a fraction of STm-infected cells initiate a partial anti-inflammatory response, but that this does not extend downstream to the expression of key anti-inflammatory genes such as *Arg1* and *Mrc1*. This suggests that *Il4ra* upregulation is insufficient to trigger full anti-inflammatory polarization.

### *Salmonella* sensitizes macrophages for an enhanced response to anti-inflammatory ligands

During infection, macrophages can be exposed to both pro- and anti-inflammatory signals, as immune cell recruitment to the site of infection releases anti-inflammatory cytokines to resolve inflammation. Upregulating IL-4Rɑ on the surface of infected macrophages may enable STm to respond to IL-4 released into the environment, thereby creating an anti-inflammatory cell state more permissive for bacterial replication. To evaluate whether there are functional consequences to the infection-driven upregulation of IL-4Rɑ, we used smFISH to compare *Arg1* expression in cells treated with M2 ligands alone, STm alone, or STm in combination with M2 ligands. At 4 hours, *Arg1* expression remained low across single and dual-stimulus treatments (Figure S3.3). However, at 24 hours, the combined treatment of STm and M2 ligands significantly increased *Arg1* expression in STm-infected and uninfected cells (Figure 3C, Table S1). The fraction of cells expressing *Arg1* was higher in infected than uninfected cells (74.8% vs 61.2%), as were overall *Arg1* expression levels per cell (Table S1). Despite increases in surface IL-4Rɑ being primarily restricted to infected cells (Figure 3B), the fraction of uninfected cells inducing *Arg1* expression in response to M2 ligands also increased compared to M2 ligands alone (61.2% vs 49.9%) (Figure 3C). Together, these data demonstrate that STm sensitizes macrophages for an enhanced anti-inflammatory response, and while IL-4 acts as the key trigger to transition from a partial to a full anti-inflammatory response, this STm-based sensitization does not require IL-4Rɑ upregulation.

### Anti-inflammatory gene expression scales with bacterial load

Our results indicate that this enhanced *Arg1* induction in response to dual stimulation with STm and M2 ligands is largely independent of IL-4Rɑ upregulation. However, while the STm-infected population had increased *Arg1* expression compared to the uninfected population, the response remained highly heterogeneous (Figure 3C, Table S1). This suggests that the increase in surface IL-4Rɑ primarily detected in a subset of infected cells (Figure 3B) may further amplify the magnitude of the *Arg1* expression response. Previous studies have linked BMDMs containing replicating STm to increases in surface IL-4Rɑ (Saliba *et al*., 2016; Stapels *et al*., 2018), and gene expression patterns consistent with M2 polarization have been shown to correlate with increasing bacterial load (Heyman *et al*., 2023). Therefore, we hypothesized that bacterial load is responsible for the variation in anti-inflammatory gene expression we observe in infected cells.

To test this, we used smFISH to compare *Il4ra* and *Arg1* gene expression at 24 hours in response to STm, with macrophages classified into four subpopulations according to bacterial load (uninfected, low, medium, high; see Methods, Figure S4.1). For *Il4ra*, we examined cells exposed to STm alone, while for *Arg1*, we examined cells exposed to STm and M2 ligands, as infection alone does not induce *Arg1* expression (Figure 2A). To quantify changes in expression, we defined a basal expression threshold using untreated cells and we identified high expressing cells as those above the 95^th^ percentile of expression across the pooled populations (see Methods). For both genes, expression increased with bacterial load (Figure 4A, Table S1). *Il4ra* expression was distinct in cells with low loads of STm compared to uninfected cells, and progressively increased in cells with medium and high bacterial loads. Cells with high loads of STm contained the largest fraction of *Il4ra*-high cells (mean = 28%). These results are consistent with our flow cytometry data (Figure 3B), but provide finer resolution of the relationship between bacterial load and *Il4ra* expression. Similar trends were observed for *Arg1* in the dual stimulus condition, with *Arg1*-high cells enriched among macrophages with medium to high bacterial loads (Figure 4A, Table S1). However, uninfected cells and cells with a low load of STm had similar expression profiles, indicating that increased bacterial loads may be required for high *Arg1* expression in the dual stimulus condition. These data demonstrate that *Il4ra* and *Arg1* expression scale with bacterial load. This dose-dependent relationship suggests that the upregulation of *Il4ra* in cells with high bacterial loads amplifies *Arg1* gene expression in the dual stimulus condition, linking bacterial loads to increased anti-inflammatory sensitization.

**Figure 4.**
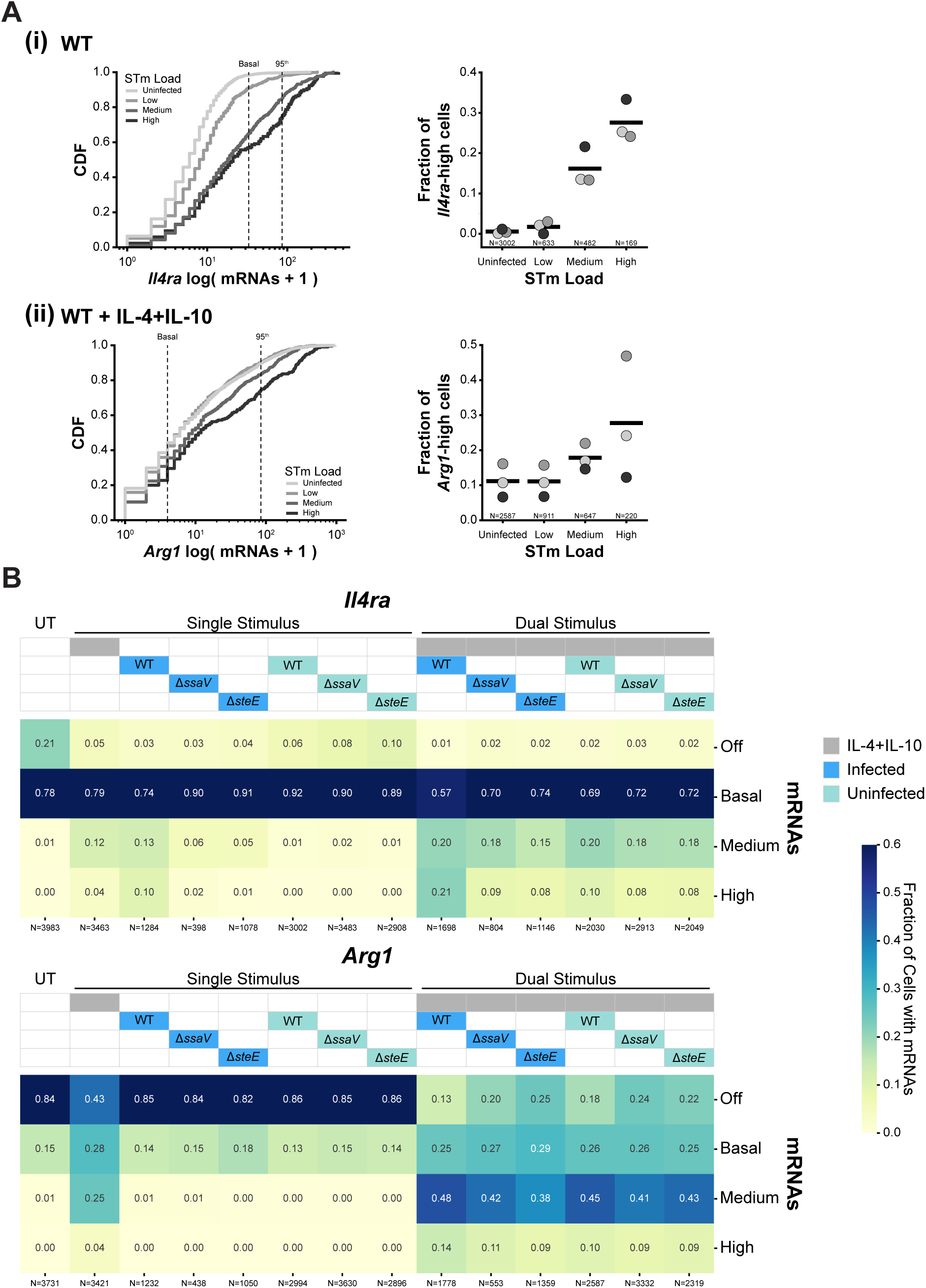
Features of infection alter the magnitude of anti-inflammatory gene expression responses. (A) Anti-inflammatory gene expression scales with bacterial load in infected CIMs. Gene expression was measured using smFISH at 24 hours, and STm-exposed CIMs classified based on their STm load (see Figure S4.1 and Methods). (i) *Il4ra* expression was measured in CIMs exposed to WT STm. (ii) *Arg1* expression was measured in CIMs exposed to both WT STm and M2 ligands (25 ng/mL IL-4 and 25 ng/mL IL-10). For each subpanel: (Left) The cumulative distribution function (CDF), plotted as log(mRNA +1) for each gene, and pooled from three biological replicates. Different STm loads are shown in shades of gray. Dashed lines indicate basal expression (32 mRNAs for *Il4ra* and 3 mRNAs for *Arg1*) and the 95^th^ percentile threshold (85 mRNAs for *Il4ra* and 84 mRNAs for *Arg1*) defining high-expressing CIMs. (Right) The fraction of high-expressing CIMs (above the 95^th^ percentile) in each STm load category. Biological replicates are shown in shades of gray with the mean shown as a black bar. N=total number of CIMs. p<0.001 for all pairwise comparisons except for *Il4ra*: medium vs high (p<0.01) and *Arg1*: medium vs high (p<0.01), uninfected vs low (p>0.05, not significant). Pairwise comparisons used a binomial generalized linear model (GLM) with post-hoc pairwise comparisons performed using linear contrasts on the fitted model coefficients, with Wald tests used to evaluate significance, with FDR correction. Results of statistical analyses on the fraction of *Il4ra*-high and *Arg1*-high across different bacterial loads are reported in Table S1. (B) *Il4ra* and *Arg1* expression are differentially dependent on STm effector secretion. CIMs were left untreated (UT), treated with M2 ligands, or exposed to STm (WT, Δ*ssaV*, or Δ*steE*) either alone (single stimulus) or in combination with M2 ligands (dual stimulus). *Il4ra* and *Arg1* expression were measured using smFISH at 24 hours. CIMs were classified as infected or uninfected as described in Figure 1 (see Methods). CIMs were binned into four expression categories based on mRNA counts using the thresholds defined in (A): Off (0 mRNAs), basal (1 to basal threshold), medium (basal threshold to 95^th^ percentile), and high (above the 95^th^ percentile). The fraction of CIMs in each category across all conditions is shown as a heatmap. Heatmap color scale is capped at 0.6 to better visualize differences between categories containing low fractions of CIMs. Data from (A) are included in the heatmaps. Data are pooled from three biological replicates. All STm infections were done at MOI 10. N=total number of CIMs.

### *Il4ra* and *Arg1* expression are differentially dependent on STm effector secretion

The dose-dependent relationship between anti-inflammatory gene expression and bacterial load could arise as a direct result of bacterial replication or alternatively, through SPI-2 effector secretion that accompanies this replication. Assembly of the SPI-2 T3SS is essential for intracellular replication (Cirillo *et al*., 1998), and recent studies have connected the SPI-2 effector SteE to IL-4Rɑ upregulation (Saliba *et al*., 2016; Jaslow *et al*., 2018; Stapels *et al*., 2018; Gibbs *et al*., 2020; Panagi *et al*., 2020). To determine the relative contributions of bacterial replication and SteE effector secretion to the increase in *IL4ra*- and *Arg1*-high cells, we used smFISH to compare *Il4ra* and *Arg1* expression at 24 hours in cells infected with WT STm, an STm mutant strain unable to assemble the SPI-2 T3SS (Δ*ssaV*), or an STm mutant strain lacking SteE (Δ*steE*). As expected, both the number of infected cells at 24 hours and intracellular bacterial replication were reduced in the Δ*ssaV* strain, while in the Δ*steE* strain, replication was more similar to WT (Figure S4.2). We classified cells into four expression bins [off (0 mRNAs), basal (1 to basal threshold), medium (basal threshold to 95^th^ percentile), and high (above the 95^th^ percentile)] using the basal and 95^th^ percentile thresholds defined in Figure 4A and compared the fraction of *IL4ra*-and *Arg1*-high cells across strains and treatments.

The fraction of *Il4ra*-high cells present in WT-infected cells was greatly reduced in both mutant strains (WT: 0.10; Δ*ssaV*: 0.02; Δ*steE*: 0.01) (Figures 4B, S4.3A). However, cells infected with the mutant strains remained distinct from uninfected cells, maintaining a small fraction of *Il4ra*-high cells (0.02 and 0.01 for Δ*ssaV* and Δ*steE,* respectively, compared to 0 for uninfected cells). This indicates that while SteE secretion drives the majority of *Il4ra*-high cells, infection alone is sufficient to induce a modest increase in these cells above baseline, independent of SteE. Next, we evaluated whether the dose-dependent relationship between bacterial load and *Il4ra* gene expression was maintained in the mutant strains. The enrichment of *Il4ra*-high cells in macrophages with high bacterial loads observed with the WT strain (Figure 4A) was lost in cells infected with either mutant strain (Figure S4.4A). However, the mutant strains differed in the scaling of gene expression with bacterial load. The Δ*steE* strain maintained the dose-dependent relationship between *Il4ra* gene expression and bacterial load, although the magnitude of the expression differences between the load categories was reduced compared to the WT strain, while the ΔssaV strain lost this relationship (Figure S4.4A). We note that substantially fewer macrophages infected with the Δ*ssaV* strain were classified as high bacterial load due to the known replication defects of this strain (Figure S4.2), which limits our ability to interpret dose-dependent relationships in this strain. Together, these data indicate that high levels of *Il4ra* upregulation are primarily dependent on SteE secretion, especially in cells with high bacterial loads, but that additional effector proteins or other T3SS-dependent processes are involved in establishing the dose-dependent relationship between bacterial load and *Il4ra* gene expression.

Given that M2 ligands induce *Il4ra* gene expression (Figure 3A), we wondered whether treatment with M2 ligands could compensate for the loss of SteE and restore the fraction of *Il4ra*-high cells to levels observed with WT infection. To test this, we compared the fraction of *Il4ra*-high cells in macrophages infected with the different STm strains and treated with M2 ligands (Figures 4B, S4.3A). While the addition of M2 ligands enhanced *Il4ra* expression and increased the fraction of *Il4ra*-high cells across all strains, WT STm consistently generated the highest fraction of these cells (0.21). In the dual-stimulus condition, the fraction of *Il4ra*-high cells in macrophages infected with the mutant strains (0.09 for Δ*ssaV*, 0.08 for Δ*steE*) was similar to uninfected cells (0.08 for both Δ*ssaV* and Δ*steE*), indicating that the addition of ligands cannot compensate for the loss of effector-mediated *Il4ra*-upregulation. Furthermore, M2 ligands differentially affected the scaling between bacterial load and *Il4ra* expression in WT and mutant infections (Figure S4.4B). Cells infected with the WT strain maintained the scaling relationship in the dual-stimulus condition; however, only cells with medium and high loads were distinct, while uninfected cells and those with low STm loads now had similar expression levels. In contrast, in cells infected with the Δ*steE* strain, the addition of M2 ligands collapsed the dose-dependent relationship between bacterial load and *Il4ra* expression. Similar results were observed with Δ*ssaV*, with the caveat that cell counts were low in the high bacterial load category. Together, these data demonstrate that effectors play critical roles in *Il4ra* upregulation that cannot be fully compensated for by the addition of M2 ligands. While environmental M2 ligands dominate the response in cells infected with effector-deficient STm or those with low bacterial loads, the effector-mediated upregulation of *Il4ra* is maintained in cells with high bacterial loads, even in the presence of exogenous ligands.

In contrast to *Il4ra*, the fraction of *Arg1*-high cells in dual-stimulus conditions was largely independent of effector secretion (Figures 4B, S4.3B). All STm strains induced similar fractions of *Arg1*-high cells (0.09-0.14), with only modest differences between infected and uninfected cells. However, the scaling of *Arg1* expression with bacterial load in WT-infected cells exposed to M2 ligands was lost in both mutant strains (Figure S4.4C), with all categories resembling uninfected cells exposed to M2 ligands, independent of bacterial load. This indicates that while the dose-dependent relationship between bacterial load and *Arg1* expression depends on effector secretion, robust *Arg1* induction in response to STm and M2 ligands does not require high bacterial loads or effector secretion.

Taken together, these results demonstrate that fundamentally different mechanisms are responsible for inducing *Il4ra-* and *Arg1-high* cells during STm infection. *Il4ra*-high cells depend on SteE secretion, and the role of effectors in *Il4ra* upregulation cannot be fully rescued by the addition of exogenous M2 ligands. In contrast, *Arg1*-high cells only require STm exposure followed by M2 ligands, with effector secretion having negligible effects. Furthermore, the consistent loss of scaling of gene expression with bacterial load in macrophages infected with the mutant strains and treated with M2 ligands suggests that effector secretion or bacterial replication amplifies differences in anti-inflammatory gene expression in infected cells.

## DISCUSSION

Here, we characterized the signals required for anti-inflammatory polarization of macrophages during STm infection of CIMs. Using smFISH, we show that STm fails to robustly induce pro- or anti-inflammatory responses. Instead, we find that STm establishes a sensitized cell state characterized by enhanced responsiveness to the anti-inflammatory cytokine IL-4. Exposing cells to both STm and IL-4 generates an amplified anti-inflammatory gene expression response, particularly for *Arg1*. We find that this sensitization occurs largely independent of IL-4Rɑ surface upregulation, with STm exposure sufficient to trigger the enhanced cytokine responsiveness. However, we demonstrate that features of the infection state, including bacterial load and effector secretion, can modulate the magnitude of the polarization response, with high bacterial loads associated with SteE-dependent *Il4ra* upregulation. These context-dependent responses generate a spectrum of anti-inflammatory states that scale with infection progression. Together, our data establish a two-signal model for macrophage polarization during infection, requiring STm-induced sensitization and environmental IL-4. Instead of directly inducing an anti-inflammatory state, STm, via unknown mechanisms, enhances macrophage responsiveness to IL-4, facilitating the integration of environmental cues into the modulation of polarization during infection.

A critical element of our two-signal model is that STm establishes a macrophage state poised for, rather than committed to, anti-inflammatory responses. While the molecular mechanisms responsible for this sensitized state remain unknown, studies of macrophage priming and M1-M2 transitions provide several possible explanations. First, although TLR signaling typically induces pro-inflammatory responses, it has also been linked to anti-inflammatory gene expression in several contexts. In mycobacteria-infected macrophages, TLR signaling through the adaptor MyD88 is required for *Arg1* expression (El Kasmi *et al*., 2008; Qualls *et al*., 2010). Prior exposure of macrophages to LPS and IFN-γ primes cells for an enhanced response to IL-4 via upregulation of several anti-inflammatory genes (O’Brien and Spiller, 2022). These results demonstrate that detection of bacteria and subsequent pro-inflammatory signaling can elicit a macrophage state primed for anti-inflammatory gene expression. Second, STm exposure could remodel chromatin, increasing accessibility of anti-inflammatory gene promoters to transcription factors such as STAT6. Our finding that *Arg1* is minimally induced by M2 ligands alone despite robust activation of STAT6 suggests that chromatin accessibility may be a limiting factor for anti-inflammatory gene expression in CIMs. Chromatin accessibility of *Arg1* has been shown to be decreased by TNF, which has also been found to limit M2 polarization in STm granulomas in mice (Schleicher *et al*., 2016; Pham *et al*., 2020). However, given that STm robustly induces *Tnf* expression in both infected and uninfected cells in our system, *Tnf* expression levels are unlikely to play a role in establishing the sensitized state. A key finding in our study is that sensitization occurs independently of IL-4Rɑ upregulation, as uninfected cells, which lack this receptor upregulation, display increased responsiveness to IL-4. This IL-4Rɑ independent mechanism may enable a broader spectrum of sensitized states that can be tuned by infection context. By separating the initial sensitization from the receptor-mediated amplification of the response, this may enable macrophages to fine-tune their responses and adjust their polarization state according to environmental signals. Identifying the molecular mechanisms responsible for this IL-4Rɑ independent mechanism will be critical to understand how pathogen exposure reprograms macrophage cell state.

Our finding that IL-4 is required for robust anti-inflammatory gene expression in STm-infected macrophages suggests that polarization is gated by the availability of environmental signals. This two-signal requirement (STm sensitization and environmental IL-4) has several implications for the pathogen. First, environmental gating may provide flexibility in STm-induced macrophage polarization responses. Infected macrophages in IL-4 replete environments, such as regions involved in tissue repair or helminth co-infections (Murray, 2017), would rapidly polarize to M2 states, whereas those in IL-4 poor environments would remain pro-inflammatory. In addition, heterogeneity in cytokine production can generate spatial diversity in cytokine environments even within a single infection site (Pham *et al*., 2020; Petrucciani *et al*., 2024; Russell *et al*., 2025). The requirement for a second signal would enable STm-infected macrophages to adapt to changing cytokine environments during infection and diversify host responses. Second, this type of two-signal gating may help avoid inappropriate polarization responses. AND logic gates are a common feature of cell fate specification during development (Liem *et al*., 2000; Voas and Rebay, 2004; ten Berge *et al*., 2008; Li and Elowitz, 2019). The requirement for coincident signal detection before committing to a developmental transition helps limit cell fate errors due to transient exposure to a single signal. Macrophage polarization is a reversible process that requires constant ligand exposure to maintain the polarized state (Liu *et al*., 2020). As such, by requiring both STm exposure and environmental IL-4, the pathogen may limit aberrant M2 polarization in response to weak or stochastic fluctuations in cytokines that may be insufficient to sustain the M2 polarized state for a prolonged period.

Our results demonstrate that anti-inflammatory polarization of infected macrophages is context-dependent, integrating features of infection to adjust the macrophage response. While all STm-exposed cells are sensitized to IL-4, the magnitude of the response varies with bacterial load and effector secretion. High bacterial loads correlate with increased *Il4ra* expression, likely driven by increased SteE secretion into the host cytoplasm. Although this *Il4ra* upregulation is not required for sensitization, receptor upregulation amplifies the cytokine response. While our experiments used saturating concentrations of IL-4, STm-infected macrophages in tissues are likely to encounter lower levels of the cytokine. By scaling the upregulation of *Il4ra* to bacterial load, highly infected macrophages may increase their sensitivity to IL-4, enabling anti-inflammatory polarization even in limiting cytokine environments and further supporting replication. Integrating bacterial features into the polarization response in this way allows for a graded rather than a binary response of STm-exposed cells to IL-4. Given the variation in bacterial state within infected macrophages and how this changes over the course of infection, a polarization response to environmental IL-4 that scales with features of infection state may be preferable to a uniform polarization response.

Anti-inflammatory polarization, *Il4ra* upregulation, and the role of SteE in receptor upregulation are known features of STm-infected macrophages (Saliba *et al*., 2016; Stapels *et al*., 2018; Gibbs *et al*., 2020; Panagi *et al*., 2020). Our work confirms these observations in CIMs, yet is also distinct, as we find that STm-mediated *Il4ra* upregulation is a mechanism that amplifies the anti-inflammatory response to subsequent environmental signals. We demonstrate that the transition to an anti-inflammatory state is a two-step process in CIMs, requiring STm-mediated sensitization and environmental IL-4. Several studies using BMDMs have reported anti-inflammatory polarization of STm-infected macrophages in the absence of exogenous IL-4 (Saliba *et al*., 2016; Stapels *et al*., 2018). This discrepancy could reflect differences between CIMs and BMDMs, such as in autocrine IL-4 production or basal IL-4Rɑ expression levels. Future comparative studies could harness these differences to further dissect mechanisms of M2 polarization. However, many features of the macrophage response to STm are consistent across the cell types, including IL-4Rɑ upregulation, the role of SteE, and the muted pro-inflammatory response, suggesting that CIMs recapitulate key aspects of STm-macrophage interactions. CIMs as a model system for STm infection offer several advantages that complement studies in primary macrophages. CIMs can be stably transduced, and permanent stocks of progenitor cells can be generated, enabling future experiments using signaling reporters and genetic perturbations to further characterize heterogeneity in macrophage responses to STm. In addition, the requirement for IL-4 in CIMs separates the sensitization and polarization steps, which will facilitate mechanistic dissection of this two-step process.

In summary, our work characterizes heterogeneity in macrophage anti-inflammatory polarization during STm infection and shows that macrophages integrate environmental cues and multiple features of infection state to generate context-dependent anti-inflammatory responses. Many other intracellular pathogens can replicate within macrophages; as such, this two-signal, context-dependent model of anti-inflammatory polarization may be a common strategy among these pathogens.

## ACKNOWLEDGEMENTS

We thank all the members of the Lane Lab for their valuable feedback and thoughtful discussions in performing this work and preparing this manuscript. Flow cytometry equipment and training were provided by the Flow Cytometry Core Facility in the Robert H. Lurie Comprehensive Cancer Center at Northwestern University (NCI Cancer Center Support Grant #P30 CA060553) and the Single Cell Genomics Facility at Northwestern University (RRID:SCR_026652). GA was supported by the Cellular and Molecular Basis of Disease Training Program (NIH T32 GM008061).

## MATERIALS and METHODS

### Mammalian Cell Culture

Hoxb8 conditionally immortalized macrophage (CIM) progenitors, B16-GM-CSF, and 3T3-M-CSF cells were obtained from the Cox lab (Roberts *et al*., 2019). Media formulations: 1) CIM progenitor media: RPMI 1640 medium (Fisher Scientific #11875119) supplemented with 10% fetal bovine serum (FBS) (Omega Scientific #FB-02), 2 mM GlutaMAX (Thermo Fisher Scientific #35050061), 1 mM sodium pyruvate (Fisher Scientific #SH3023901, Sigma Aldrich #S8636), 10 mM HEPES buffer (Fisher Scientific #15630080), 10 µg/mL penicillin-streptomycin solution (Fisher Scientific #MT30002CI), 43 µM ꞵ-mercaptoethanol (Sigma Aldrich #M3148), 2% GM-CSF conditioned media (prepared from B16-GM-CSF cells), and 2 µM ꞵ-estradiol (Sigma Aldrich #E2758). 2) Macrophage-differentiation media: DMEM (Fisher Scientific #MT15017CV) supplemented with 10% FBS, 2 mM GlutaMAX, 10 µg/mL penicillin-streptomycin solution, 1 mM sodium pyruvate, and 10% M-CSF conditioned media (prepared from 3T3-M-CSF cells). Efficiency of each batch of M-CSF conditioned media was evaluated by measuring the macrophage cell surface marker F4/80 (BioLegend #123110) using flow cytometry.

CIM progenitors were cultured in suspension in non-tissue culture treated flasks (Thermo Fisher Scientific #156800, #169900) at 37°C, 5% CO_2_. Cells were passaged daily by diluting to a final concentration of 1.25×10^5^ cells/mL, and density did not exceed 5×10^5^ cells/mL before passaging. To differentiate CIM progenitors, the cells were washed twice in phosphate buffered saline (PBS) (Fisher Scientific #MT21040CV) containing 1% FBS, resuspended in differentiation media and plated at 3.5×10^5^ cells per well in non-tissue culture treated 6-well plates (Thermo Fisher Scientific #150239; Genesee Scientific #25-100). 1 mL of fresh differentiation media was added to each well on days 3 and 6. Differentiated macrophages (CIMs) were harvested on day 7 for all experiments. All experiments used CIMs below passage 30, and biological triplicates were derived from at least two independent CIM progenitor stocks.

### *Salmonella* Cell Culture

The wild type (WT) *Salmonella enterica* serovar Typhimurium (STm) 14028 strain used in this study was a gift from the lab of Michael McLelland. Mutant STm strains (Δ*ssaV* and Δ*steE*) were from the single-gene mutation library (Porwollik *et al*., 2014), carry a chloramphenicol resistance cassette, and were PCR confirmed. All STm strains were made electrocompetent using the mannitol-glycerol preparation method (Warren, 2011). Plasmids constitutively expressing mCerulean3 (imaging: pTU1-A-att-SJM906-PET-mCerulean3o-Bba_B0015-att-pSC101-carb, mCerulean3 is codon optimized for STm; flow cytometry: pFPV-mCerulean3-o (Lane *et al*., 2019)) were used to transform these strains. A high copy plasmid was used for flow cytometry due to the reduced sensitivity of this technique compared to microscopy. WT STm transformants were selected on LB agar plates with carbenicillin (25 µg/mL for pTU1, 50 µg/mL for pFPV) (Thermo Fisher Scientific #50213248). Transformants from mutant STm strains were selected on LB agar plates with carbenicillin (25 µg/mL) and chloramphenicol (10 µg/mL) (Thermo Fisher Scientific #BP904-100). Strains were streaked from glycerol stocks onto LB agar plates containing the corresponding antibiotic(s) at the indicated concentration and incubated at 37°C overnight. Plates were discarded after one week.

### Macrophage Infections and Ligand Treatments

CIMs were plated the day prior to infection at 2×10^4^ cells per well on fibronectin-coated (Sigma Aldrich # F0895) glass bottom 96-well plates (Fisher Scientific #NC0536760) for imaging experiments, or at 1×10^6^ cells per well in non-tissue culture treated 6-well plates for flow cytometry experiments. STm cultures were grown overnight (14-16 hours) in LB supplemented with the corresponding antibiotic(s) at the indicated concentrations at 37°C with shaking.

One hour before infection, macrophage media was replaced with imaging media (IM) consisting of FluoroBrite DMEM (Fisher Scientific #A1896701) supplemented with 10 mM HEPES buffer, 1% FBS, and 2 mM GlutaMAX. STm were rinsed twice in PBS and diluted to a multiplicity of infection (MOI) of 10. This bacterial suspension (5 µL for imaging experiments and 50 µL for flow cytometry experiments) was added to CIMs, and plates were centrifuged at 1000 rpm for 15 minutes at 34°C to synchronize infection. Plates were returned to the incubator for 30 minutes to allow infection to occur. Following infection, CIMs were washed twice with IM to remove extracellular bacteria and maintained in IM supplemented with 10 µg/mL gentamicin for the remainder of the experiment. For ligand treatment conditions, ligands were added at this time, either 100 ng/mL LPS (Fisher Scientific #NC1198692) and 5 ng/mL IFN-γ (BioLegend #575302) for M1 ligands, or 25 ng/mL IL-4 (BioLegend #574302) and 25 ng/mL IL-10 (BioLegend #575802) for M2 ligands. CIMs were fixed with 4% paraformaldehyde (Electron Microscopy Sciences #15710) for 15 minutes at room temperature, washed with PBS, and stored in PBS at 4°C for up to 5 days before processing for smFISH or immunofluorescence.

### Single molecule fluorescent *in situ* hybridization (smFISH)

To enable segmentation of the CIM cytoplasm, CIMs were first stained with 50 µL of CellMask Green (Thermo Fisher Scientific #C37608) at room temperature for 10 minutes. The ViewRNA ISH Cell Assay Kit (Thermo Fisher Scientific #QVC0001) was then used following the manufacturer’s instructions, with modifications for 96-well glass-bottom plates. Upon completion of the protocol and staining of the nuclei with DAPI, cells were rinsed 3 times with PBS and stored in PBS at 4°C until imaging. All smFISH experiments were imaged within 24 hours of staining. ViewRNA probe sets labeled with Alexa Fluor 647 (Thermo Fisher Scientific #V-06) were used to detect the following transcripts: *Nos2* (ID: VB6-19580-VC), *Tnf* (ID: VB6-10973-VC), *Mrc1* (ID: VB6-18742-VC), *Arg1* (ID: VB6-18737-VC), and *Il4ra* (ID: VB6-17476-VC).

### Immunofluorescence (IF)

Antibodies used: rabbit-anti-p65 (Cell Signaling Technology (CST) #8242, 1:400), rabbit-anti-p-STAT6 (Tyr641) (CST #56554, 1:200), and goat anti-rabbit Cy5 (abcam #ab97077, 1:1000). All primary and secondary antibody dilutions were made in a blocking buffer consisting of 5% goat serum (Sigma Aldrich #G9023) in PBS immediately prior to staining.

All steps were performed at room temperature unless otherwise noted. Cells were permeabilized with 0.1% Triton-X100 (Sigma Aldrich #T8787) for 30 minutes. Cells were then rinsed once with PBS, incubated with blocking buffer for 1 hour, followed by the primary antibody overnight at 4°C with rocking. The following day, the plate was brought to room temperature, rinsed three times with PBS, and incubated with the secondary antibody for 1.5 hours. Cells were rinsed 3 times in PBS, stained with DAPI for 5 minutes, and rinsed once with PBS. With the exception of p65 IF experiments, cells were then stained with Alexa Fluor 488 Phalloidin (Thermo Fisher Scientific #A12379) for 1 hour, followed by 3 rinses with PBS. Cells were stored in PBS at 4°C until imaging. All IF experiments were imaged within 24 hours.

### Fluorescence Microscopy

Imaging was performed with a Nikon Ti2-E fluorescence microscope equipped with a Spectra III light engine, Prime BSI sCMOS camera, motorized stage, perfect focus system and controlled by Nikon Elements software. An Oko-Labs system provided temperature (37°C) and environmental (5% CO_2_) control. 16-bit images were acquired using a 40x/0.95 numerical aperture objective with 2×2 binning. Images of smFISH probed samples were acquired with a z optical spacing of 0.5 µm (11 total images). Excitation exposure times and intensities were as follows: DAPI: 150ms, 20% (smFISH, IF); CFP: 500ms, 50% (smFISH, IF); YFP: 500ms, 50% (smFISH), 500ms, 75% (p65 IF), 100ms, 45% (p-STAT6 IF); Cy5: 150ms, 25% (smFISH), 500ms, 45% (IF).

### Image Analysis

#### CIM Cytoplasm and Nucleus Segmentation

Segmentation was performed using Cellpose 2.2 with a flow threshold of 0.81 and a cell probability threshold of −1.81 (Pachitariu and Stringer, 2022). Nuclei were segmented using DAPI images and a custom Cellpose model trained on 37 ground truth images in Google Colab with a learning rate of 0.1, weight decay of 0.0001, and 500 epochs. Cytoplasm was segmented using YFP images (Cell Mask Green for smFISH experiments, Alexa Fluor 488 phalloidin for p-STAT6 IF, constitutively expressed eGFP for p65 IF (Roberts *et al*., 2019)) and either the Cellpose cyto2 model (IF) or a custom model (smFISH). The custom model was trained as described above using 50 ground truth images. All cytoplasm segmentation masks were manually inspected and corrected using the Cellpose 2.0 GUI.

#### STm Segmentation and Classification of Infected Cells

Bacterial segmentation was performed using CFP images of STm-mCerulean3. For smFISH, images were first background subtracted using the rolling ball background subtraction function in Fiji (Schindelin *et al*., 2012) with a radius of 3 pixels. Bacteria were segmented and linked to host cells using functions from CellTK (version 2.4.2) (Kudo *et al*., 2018) as previously described (Lane *et al*., 2019). Bacterial objects with a minimum area of 5 pixels were assigned to host cells using the cytoplasm masks. Objects outside of cytoplasm masks or with an area less than 5 pixels were excluded from further analysis. CIMs were classified as infected or uninfected based on the presence or absence of bacterial objects within the cytoplasm mask, respectively.

#### Classification of Infected Cells by Bacterial Load

To classify cells by bacterial load, features of intracellular STm objects, including total intensity (arbitrary units, A.U.) and total area (pixels), were extracted using the regionprops function from scikit-image (version 0.24.0) (Van Der Walt *et al*., 2014). For CIMs containing multiple bacterial objects, these objects were first counted and then relabeled as a single merged object before feature extraction. A total of 254 STm-infected CIMs were manually classified as containing low, medium, or high bacterial loads (Figure S4.1). For each CIM, the total fluorescence intensity and total area of all intracellular bacterial objects were quantified. Using the manually classified dataset, thresholds for total intensity, total area, and the number of bacterial objects were defined to classify infected CIMs into low, medium, or high STm load categories as follows: low (total area ≤ 25 pixels & total fluorescence intensity ≤ 2×10^5^ A.U. & total bacterial objects ≤ 2; medium (25 < total area < 150 pixels or total area < 150 pixels & total intensity > 2×10^5^ A.U.); high (total area ≥ 150 pixels or total intensity ≥ 1.5×10^6^ A.U.).

#### smFISH

mRNA spot detection was performed using the detect_spots function from FISH-quant-v2 (version 0.6.2) (Imbert *et al*., 2022) using a kernel size of 1.3 and a minimum distance of 1.0. For each probe, a detection threshold was set per experiment using the 10^th^ percentile of automatic thresholds generated by the get_elbow_values function for all images in a given replicate. This threshold was then used to run the detect_spots function on images from unprobed control cells. The median intensity of spots detected in these unprobed images was used as a baseline intensity threshold to identify valid spots in probed images. Valid spots were processed through the decompose_dense function to separate overlapping spots using the default alpha, beta, and gamma parameters. Cytoplasm masks were used with the extract_cell function to assign these decomposed spots to individual host cells. The infection status and mRNA spot counts for each macrophage were used for plotting and statistical analysis. For Figure 4 and related supplemental figures, basal expression thresholds for *Il4ra* and *Arg1* were defined as the 99^th^ percentile mRNA count in untreated cells (32 mRNAs for *Il4ra* and 3 mRNAs for *Arg1*). The 95^th^ percentile threshold defining *Il4ra*-high and *Arg1*-high expressing cells was determined using mRNA counts pooled from all conditions in Figure 4B (85 mRNAs for *Il4ra* and 84 mRNAs for *Arg1*). Both basal and 95^th^ percentile thresholds were calculated using pooled data from three biological replicates.

#### Immunofluorescence

Nuclear masks were eroded using the binary_erosion function from SciPy (version 1.13.1) (Virtanen *et al*., 2020). Cytoplasm rings of 5 pixels were generated from the nuclear masks using the ring_dilation_above_offset_buffer function (margin=1, buffer=2, offset=200, fill_size=50) from CellTK (version 2.4.2) (Kudo *et al*., 2018). Features of nuclear masks and cytoplasm rings were quantified using the regionprops function from scikit-image (version 0.24.0) (Virtanen *et al*., 2020). Cells with atypical morphology e.g. rounded up cells, were identified using area and roundedness metrics calculated from the cytoplasm masks. Roundness was calculated using the area and perimeter values from regionprops and the equation: 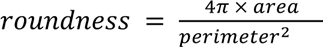. Cells with an area < 4500 pixels and a roundedness > 0.75 were excluded from further analysis. For p65, activity was calculated as the ratio of nuclear median intensity to cytoplasm ring median intensity. Baseline activity was defined as one standard deviation above the mean ratio in untreated cells. For p-STAT6, baseline activity was defined as two standard deviations above the mean nuclear median intensity in untreated cells. For both p65 and p-STAT6, these baseline activity thresholds were determined using pooled datasets for each timepoint, except for Figure 2B and Figure S2.2A where thresholds were set on a per replicate basis.

All smFISH and IF data was plotted using the Seaborn (version 0.13.0) (Waskom, 2021) and Matplotlib (version 3.9.0) (Hunter, 2007) Python packages. Python code used for image analysis, quantification of imaging data, and plotting will be available on GitHub at the time of publication.

### Flow Cytometry and Analysis

CIMs were detached using ice-cold 5 mM EDTA in PBS over ice. Cells were fixed by adding an equal volume of 8% paraformaldehyde and incubated at room temperature for 15 minutes. Fixed samples were washed twice with flow staining buffer (FSB: 5mM EDTA, 1% FBS in PBS) and stored in FSB at 4°C overnight. Samples were stained and analyzed within 1-4 days after fixation. Cells were washed once in cold FSB, resuspended at 1×10^7^ cells/mL, and incubated with TruStain FcX PLUS antibody (BioLegend #156603) following the manufacturer’s instructions. FSB was then added to adjust the concentration to 1×10^6^ cells/mL and samples were divided equally between three tubes: 1) unstained control, 2) PE rat IgG2a, κ isotype control (BioLegend #400507, 1:100), and 3) PE rat anti-mouse CD124 [IL-4Rɑ] (BD Biosciences #552509, 1:100). Cells were stained on ice for 30 minutes, washed 3 times with 10x sample volume of FSB, resuspended in FSB, and immediately analyzed on a BD LSRFortessa SORP Cell Analyzer with HTS (6-laser 18-parameter).

Data were analyzed using FlowJo (version 10.8.1). Forward scatter and side scatter gates were used to exclude debris and cell doublets before being down-sampled to 65,000 events using the FlowJo plugin DownSample (version 3.3.1). Gates for STm-infected cells (mCerulean positive) and IL-4Rɑ-PE-positive cells were set using untreated cells (Figure S3.2A). IL-4Rɑ-PE gates were set within each experiment to account for day-to-day variation in the fluorescence signal from the isotype controls (Figure S3.2A). Statistics, including the percentage of single and double-positive cells and mean fluorescence intensity (MFI), on the resulting subpopulations were exported from FlowJo. Plots were generated using the Seaborn (version 0.13.0) (Waskom, 2021) and Matplotlib (version 3.9.0) (Hunter, 2007) Python packages.

### Statistical analysis

#### smFISH spot counts across condition and time

Spot counts for a given probe were compared across condition/time pairs using linear mixed-effects models which took into account variability between replicates (3 biological replicates per experiment). Models were fit using log-transformed spot count data. Condition was a fixed effect and biological replicate a random effect. P-values were adjusted for multiple testing using the Benjamini-Hochberg FDR correction.

#### Gene expression across different bacterial loads (*Figure 4A*, S4.4)

Comparisons of fractions of *Il4ra*-high and *Arg1*-high cells across bacterial load categories (uninfected, low, medium, high) used a binomial generalized linear model (GLM), with data from three biological replicates. Post-hoc pairwise comparisons were performed using linear contrasts on the fitted model coefficients, with Wald tests used to evaluate significance. P-values were adjusted for multiple testing using the Benjamini-Hochberg FDR correction. For Figure S4.4A (Δ*ssaV*), since only two of three replicates had cells with high loads of bacteria this condition was not included in the statistical analysis.

#### Infection frequency and bacterial replication (Figure S4.2)

For both infection frequency and bacterial area measurements, comparisons across different STm strains in the presence or absence of M2 ligands were performed using Mann-Whitney U tests with data from three biological replicates per strain/condition pair. For infection frequency, the fraction of infected cells was calculated within each replicate for a given strain/condition pair. For bacterial area, the mean bacterial area of infected cells was calculated per replicate for a given strain/condition pair. Pairwise comparisons were made against the WT STm strain in the absence of M2 ligands, with p-values adjusted for multiple testing using the Benjamini-Hochberg FDR correction. Statistical analyses were performed in Python (scipy (version 1.13.1) (Virtanen *et al*., 2020), statsmodels (version 0.13.2) (Seabold and Perktold, 2010), scikit-posthocs (version 0.11.4) (Terpilowski, 2019)) and are reported in Table S1. Alpha set to 0.05. AI was used to guide the selection of appropriate statistical tests.

---

**Figure S1.1.**
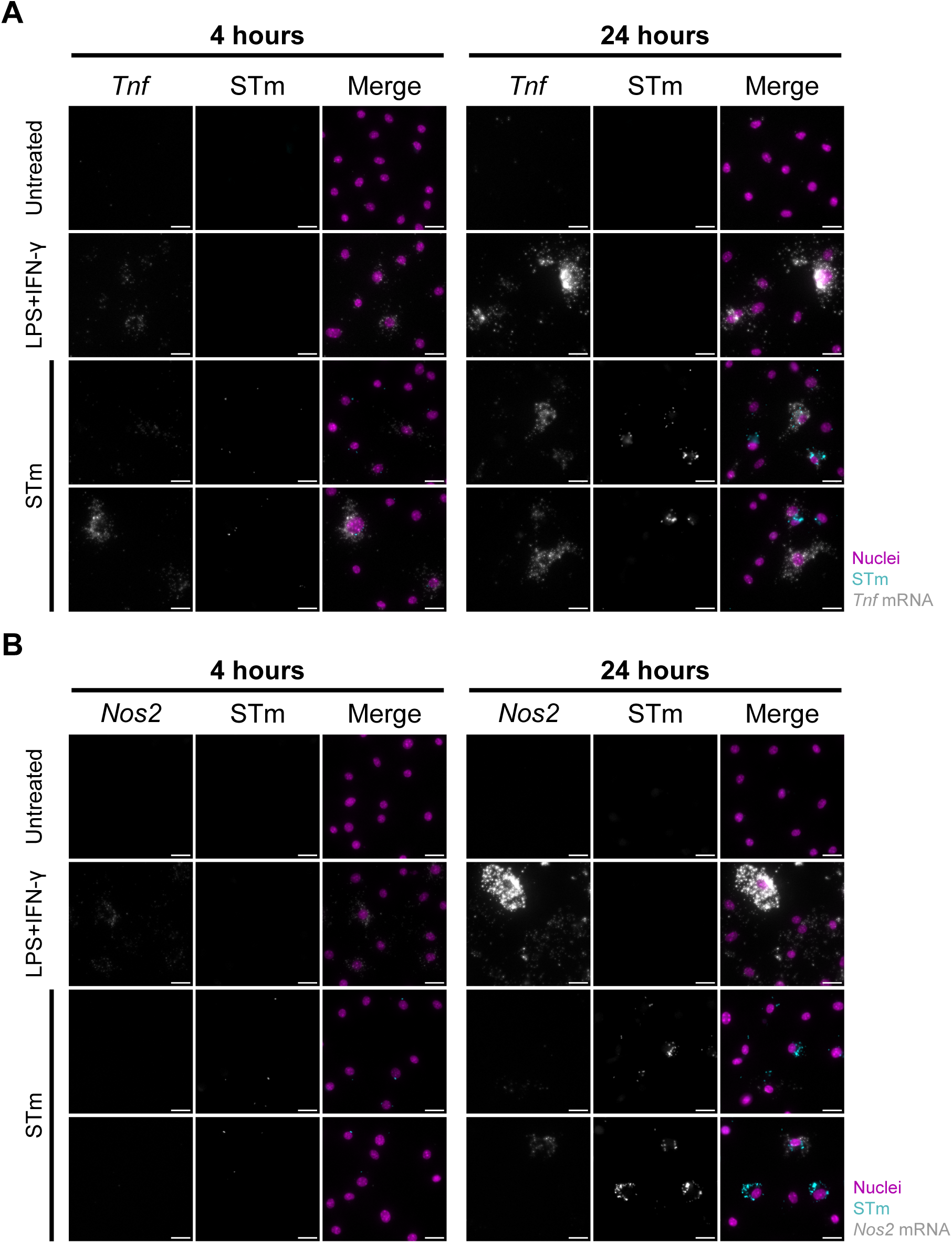
STm does not induce robust Nos2 expression. Representative smFISH images for (A) *Tnf* and (B) *Nos2* corresponding to data in Figure 1B. Images show individual channels for the smFISH probe and STm, plus merged channels (smFISH in grayscale, STm in cyan, nuclei in magenta). Scale bars=20 µm.

**Figure S1.2.**
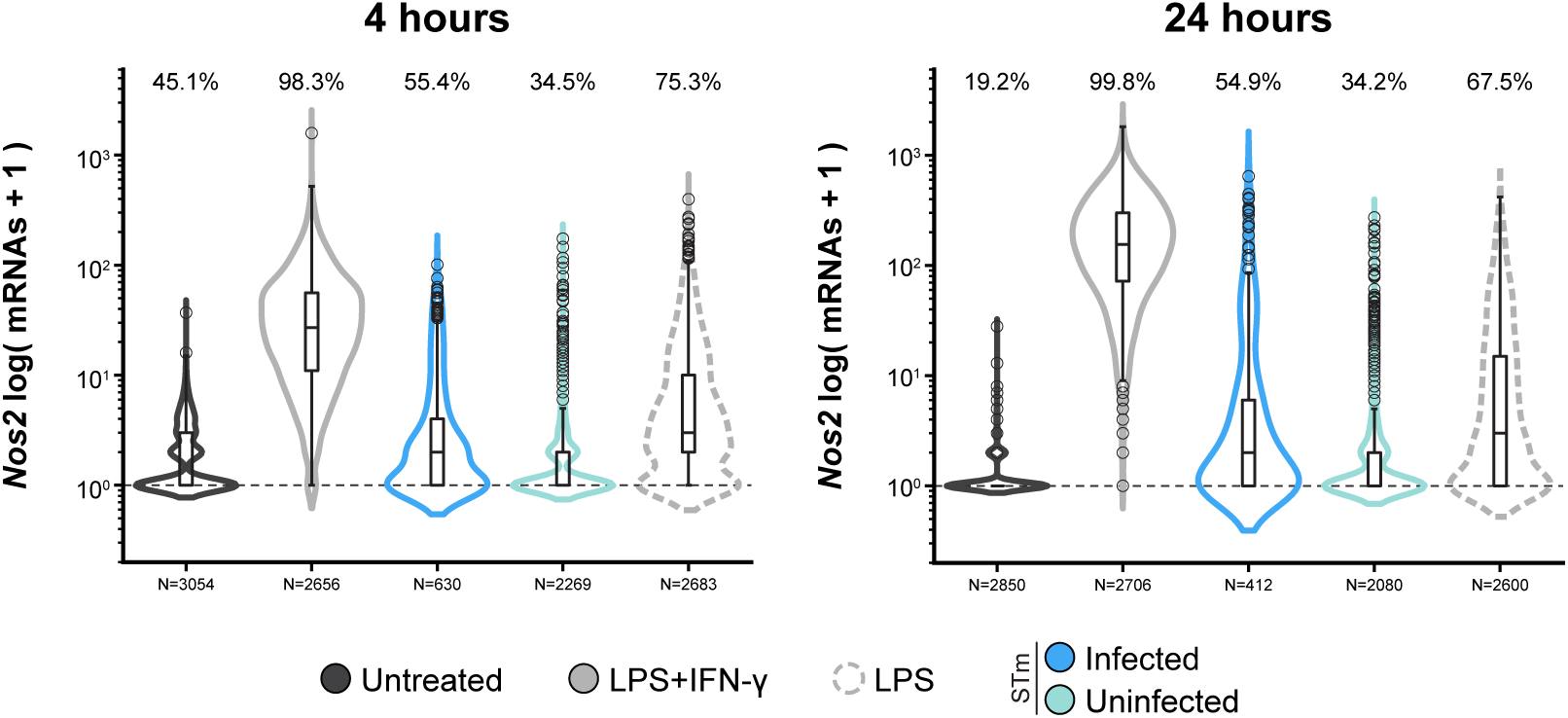
STm-induced Nos2 gene expression is comparable to LPS treatment. CIMs were left untreated, treated with LPS (100 ng/mL) alone, M1 ligands (100 ng/mL LPS and 5 ng/mL IFN-γ), or exposed to STm (MOI 10). Samples were fixed at 4 and 24 hours and *Nos2* expression was measured using smFISH. Violin plots show transcript levels; the width represents the proportion of CIMs within the respective condition. Data are presented as log(mRNA +1). Dashed line indicates expressing CIMs and the percentage of CIMs containing transcripts are above the violins. Data are pooled from three biological replicates, N=total number of CIMs. Results of statistical analyses on smFISH data are reported in Table S1.

**Figure S1.3.**
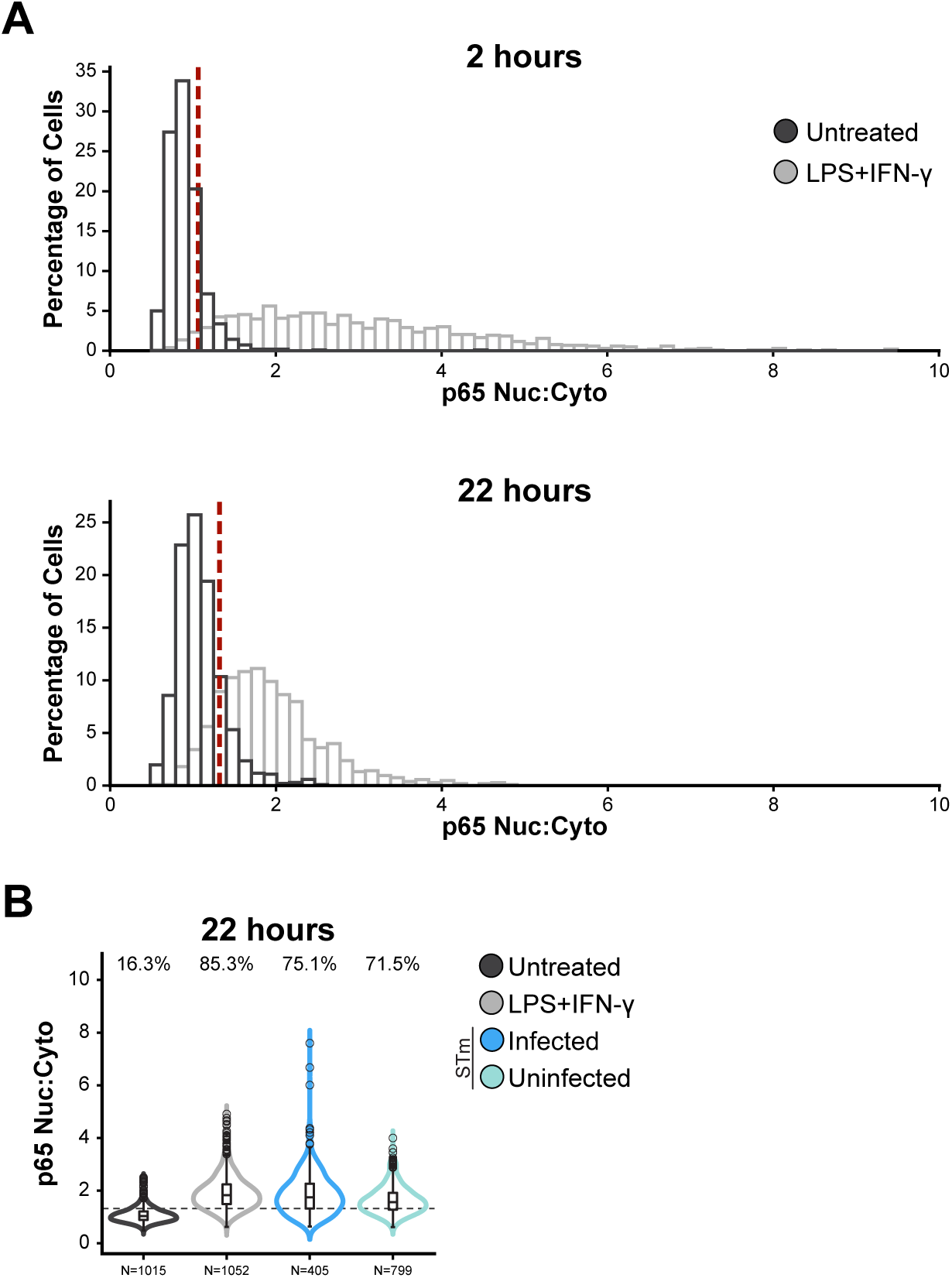
p65 signaling remains active in stimulated CIMs at 22 hours. (A) Defining the p65 basal signaling threshold used in Figure 1C. p65 localization was measured using IF at 2 and 22 hours. The p65 basal signaling threshold (red dashed line) was defined as one standard deviation above the mean p65 nuclear:cytoplasmic ratio (p65 Nuc:Cyto) of untreated CIMs at each timepoint. Histograms show the p65 Nuc:Cyto distribution in untreated (black) and LPS+IFN-γ treated (gray; 100 ng/mL LPS and 5 ng/mL IFN-γ) CIMs. Data correspond to Figures 1C and S1.3B. (B) p65 remains activated at 22 hours across all stimulated conditions. Violin plots show p65 Nuc:Cyto ratios at 22 hours measured by IF. Percentages above violins indicate the percentage of CIMs with p65 signal above the baseline (dashed line). Data are pooled from three biological replicates, N=total number of CIMs. The width of violins represents the proportion of CIMs within the respective condition.

**Figure S1.4.**
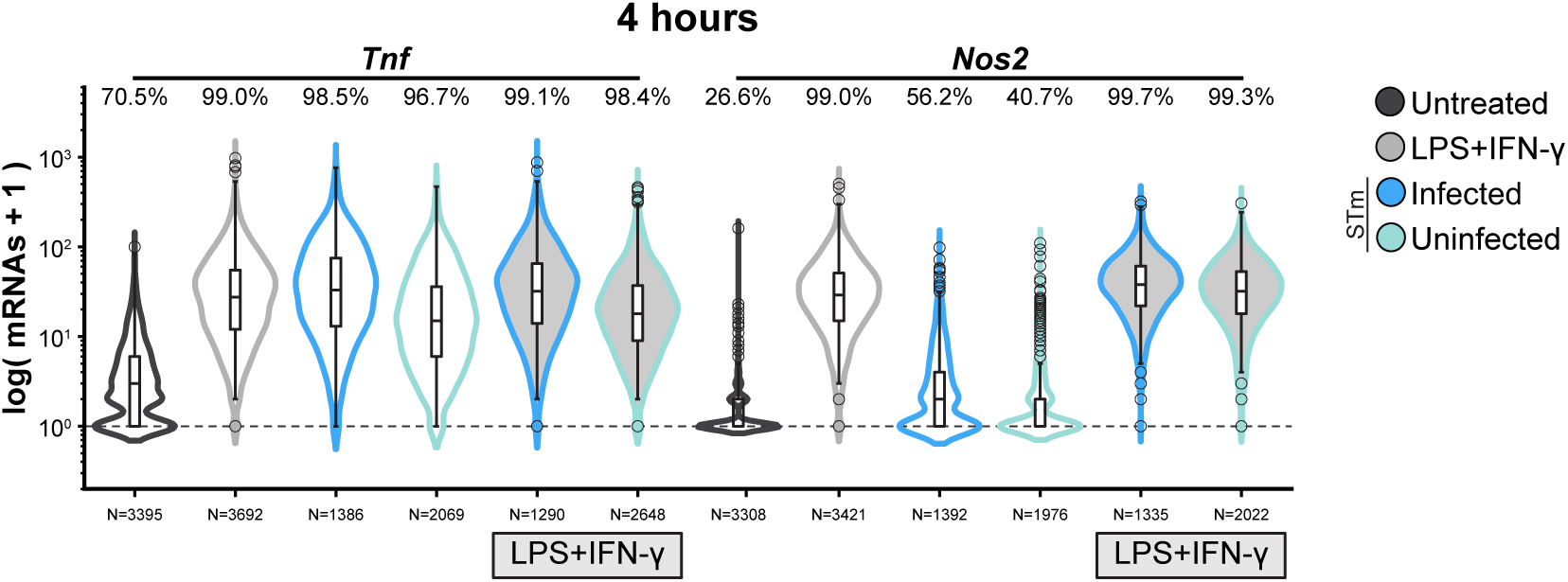
STm-exposed CIMs respond to M1 ligands at 4 hours and induce Nos2. *Tnf* and *Nos2* expression were measured at 4 hours using smFISH. CIMs were left untreated or treated with M1 ligands (100 ng/mL LPS and 5 ng/mL IFN-γ), exposed to STm (MOI 10) (empty violins), or exposed to STm (MOI 10) and treated with M1 ligands (100 ng/mL LPS and 5 ng/mL IFN-γ) (filled violins). Data are presented as log(mRNA +1). Data are pooled from three biological replicates, N=total number of CIMs. The width of violins represents the proportion of CIMs within the respective condition. Results of statistical analyses on smFISH data are reported in Table S1.

**Figure S2.1.**
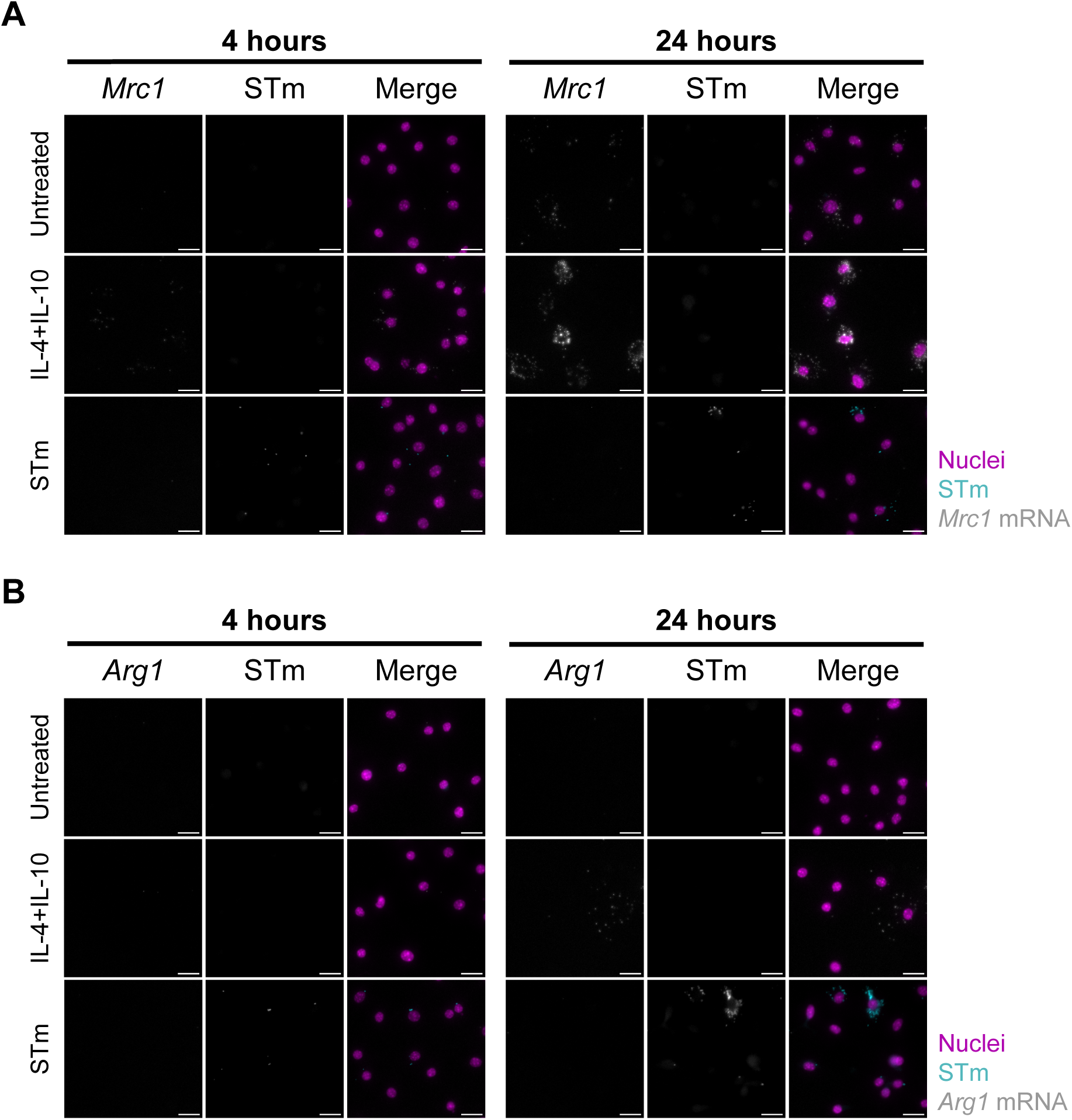
STm does not induce Arg1 or Mrc1 gene expression. Representative smFISH images for (A) *Mrc1* and (B) *Arg1* corresponding to data in Figure 2A. Images show individual channels for the smFISH probe and STm, plus merged channels (smFISH in grayscale, STm in cyan, nuclei in magenta). Scale bars=20 µm.

**Figure S2.2.**
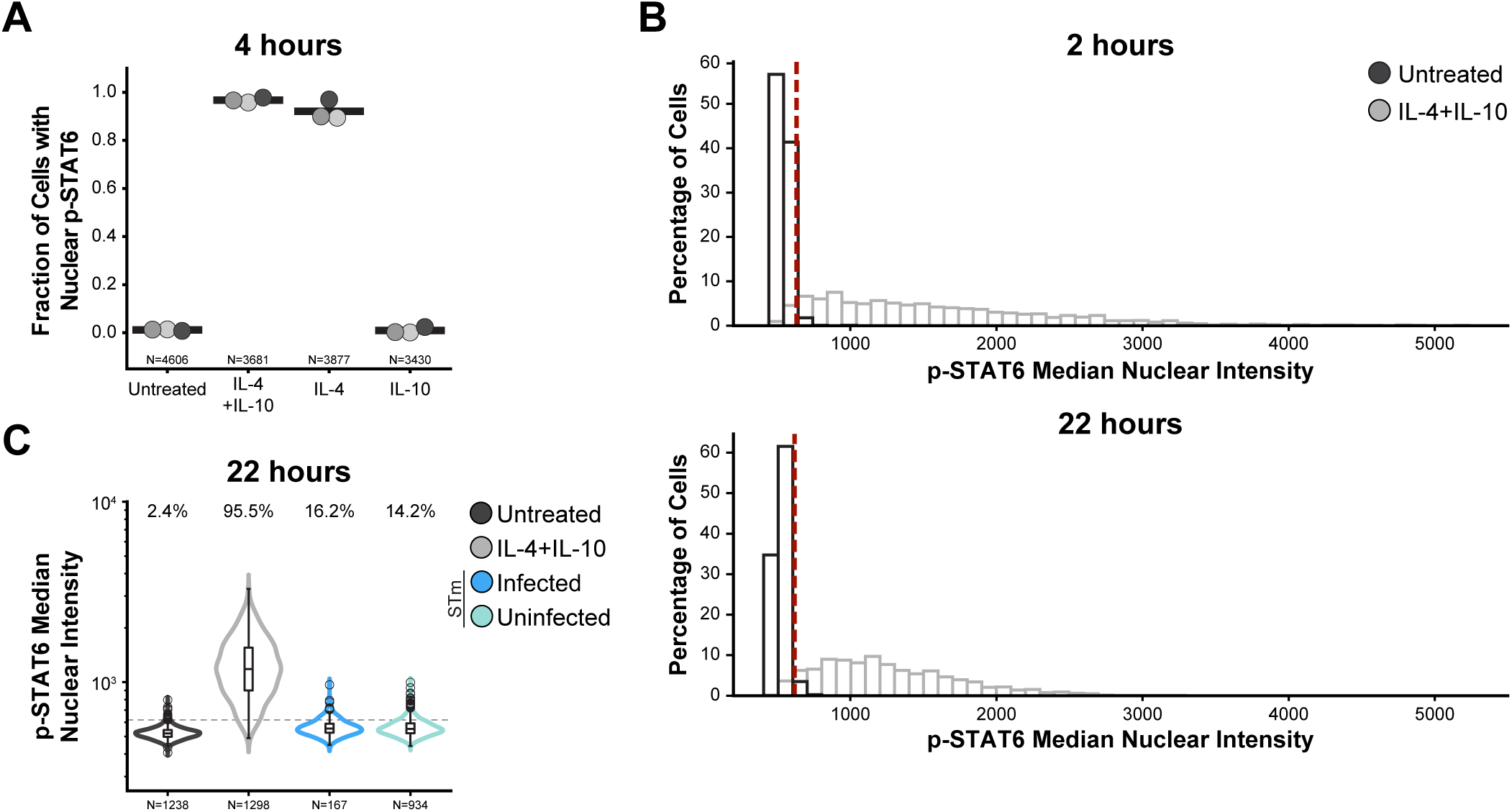
p-STAT6 signaling quantification and basal threshold definition. (A) IL-4 drives STAT6 phosphorylation in CIMs. CIMs were left untreated or treated with IL-4 (25 ng/mL), IL-10 (25 ng/mL), or both ligands, and p-STAT6 was measured by IF at 4 hours corresponding to Figure 2B. The mean fraction of CIMs with nuclear p-STAT6 above the basal signal threshold is shown for three biological replicates (shades of gray). Basal signal threshold was set per replicate (see Methods). (B) Defining the p-STAT6 basal signal threshold used in Figures 2B-C and S2.2A, C. p-STAT6 was measured by IF at 2 and 22 hours. Histograms show the median nuclear p-STAT6 intensity in CIMs that were untreated (black) or treated with M2 ligands (gray; 25 ng/mL IL-4 and 25 ng/mL IL-10). The p-STAT6 basal signal threshold (red dashed line) was defined as two standard deviations above the mean nuclear p-STAT6 signal of untreated CIMs at each timepoint. (C) STm does not induce STAT6 phosphorylation at 24 hours. CIMs were left untreated, treated with M2 ligands, or exposed to STm (MOI 10). Samples were fixed at 24 hours, stained for p-STAT6, and imaged. Violin plots show median nuclear p-STAT6 intensity; the width of violins represent the proportion of CIMs within the respective condition. Dashed line indicates basal signal threshold (defined in (B) and in Methods). Data shown in (B) and (C) are pooled from three biological replicates, N=total number of CIMs.

**Figure S3.1.**
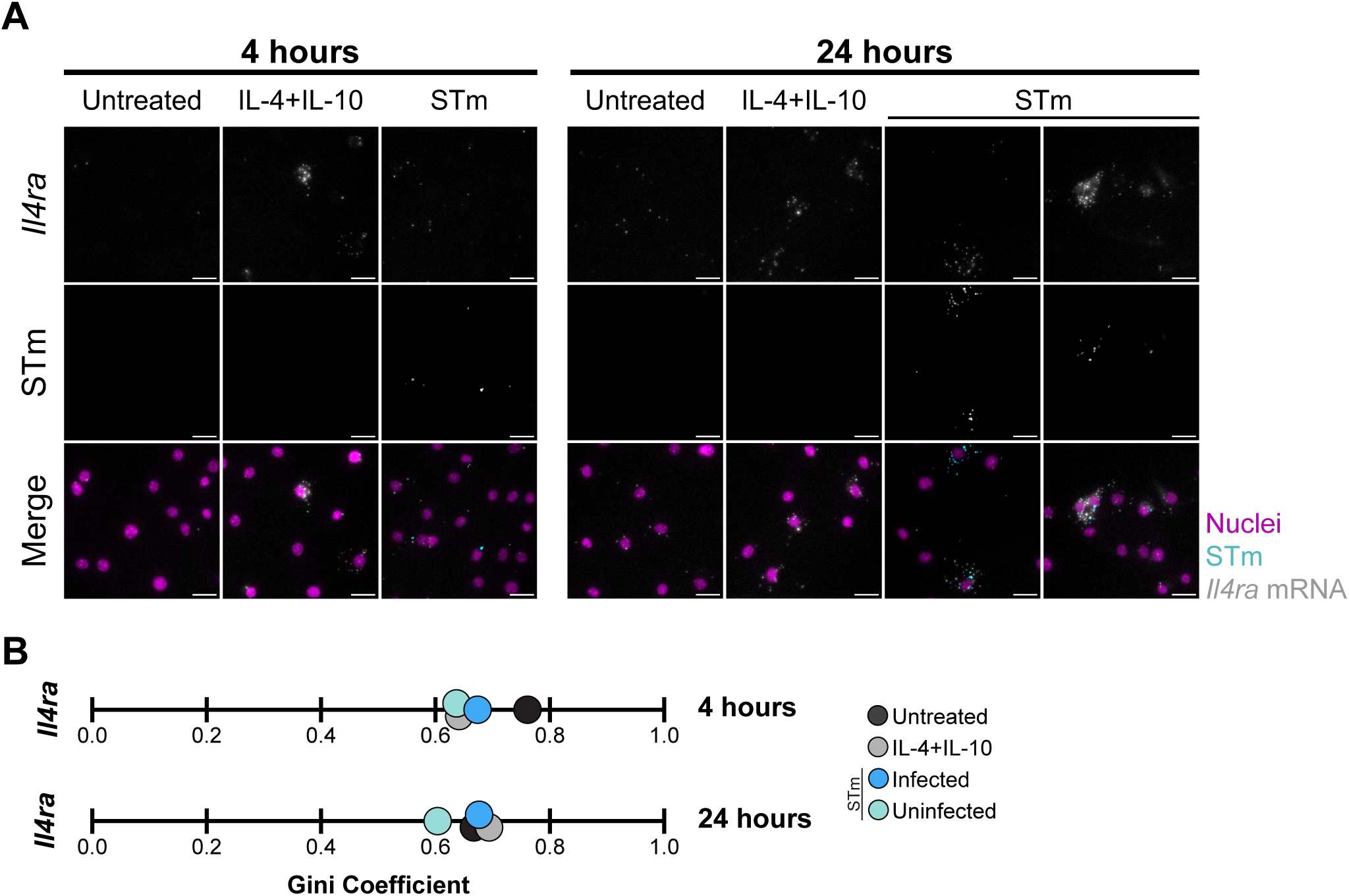
Heterogeneity in Il4ra gene expression is similar across conditions. Data corresponds to that in Figure 3A. (A) Representative smFISH images for *Il4ra*. Images show individual channels for the smFISH probe and STm, plus merged channels (smFISH in grayscale, STm in cyan, nuclei in magenta). Scale bars=20 µm. (B) Gini coefficients quantifying expression heterogeneity for *Il4ra* expression at 4 and 24 hours. Gini coefficients are reported in Table S1.

**Figure S3.2.**
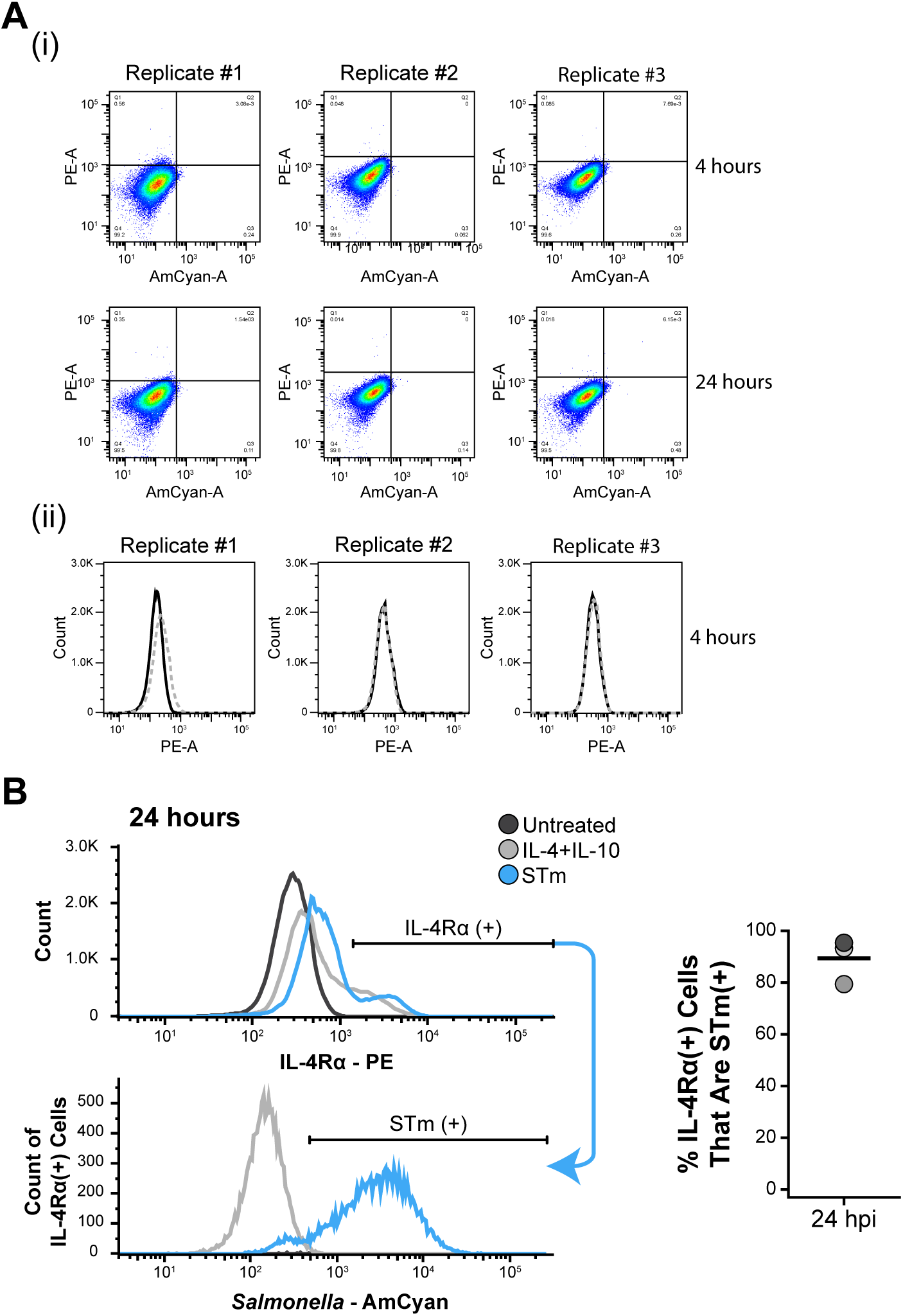
IL-4Rɑ surface protein levels increase in STm-infected CIMs. Flow cytometry analysis of IL-4Rɑ surface expression (IL-4Rɑ-PE) and STm (STm-AmCyan) corresponding to Figure 3B. (A) (i) IL-4Rɑ-PE and STm-AmCyan gating for each individual replicate. Density plots of untreated CIMs stained for IL-4Rɑ are shown at 4 and 24 hours. (ii) Isotype control signal (dashed gray line) overlaps with IL-4Rɑ-PE (solid black line) signal in untreated CIMs, with some day-to-day variation in the degree of overlap. Plots display data from three biological replicates. (B) IL-4Rɑ protein levels increase primarily in STm-infected CIMs. CIMS were left untreated (black), treated with M2 ligands (gray; 25 ng/mL IL-4 and 25 ng/mL IL-10), or exposed to STm (blue; MOI 10), fixed at 24 hours, and analyzed by flow cytometry. A representative histogram from a single experiment shows IL-4Rɑ-PE signal (top). The IL-4Rɑ+ gate was set using untreated CIMs. CIMs identified as IL-4Rɑ+ were analyzed for STm-AmCyan signal (bottom). Quantification shows the fraction of STm-exposed CIMs that are IL-4Rɑ+ and are also positive for STm. Three biological replicates (shades of gray) at 24 hours. Black bars indicate the mean. Samples in (A) and top of (B) contain 65,000 CIMs. hpi = hours post-infection.

**Figure S3.3.**
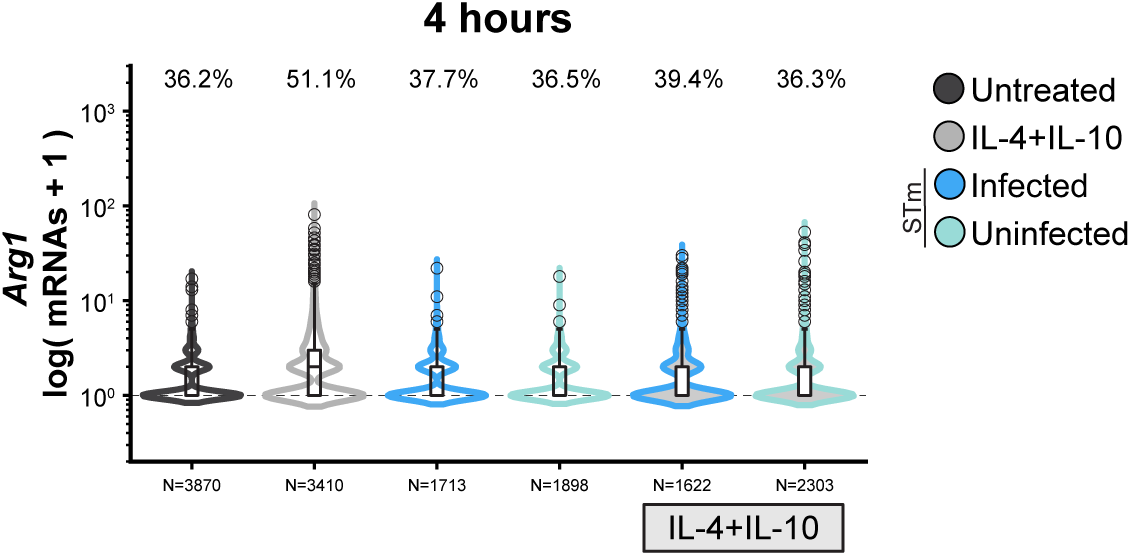
Enhanced induction of Arg1 gene expression by combined STm and M2 ligands requires extended treatment. CIMs were left untreated, treated with M2 ligands (25 ng/mL IL-4 and 25 ng/mL IL-10), exposed to STm (MOI 10) (empty violins), or exposed to STm followed by treatment with M2 ligands (filled violins). CIMs were fixed at 4 hours, processed for smFISH, and imaged. Violin plots show transcript levels for *Arg1*, data are presented as log(mRNA +1), and the width of violins represent the proportion of CIMs within the respective condition. Dashed line indicates expressing CIMs and the percentage of CIMs containing transcripts are above the violins. Data are pooled from three biological replicates, N=total number of CIMs. CIMs were classified as infected or uninfected as in Figure 1 (see Methods). Results of statistical analyses on smFISH data are reported in Table S1.

**Figure S4.1.**
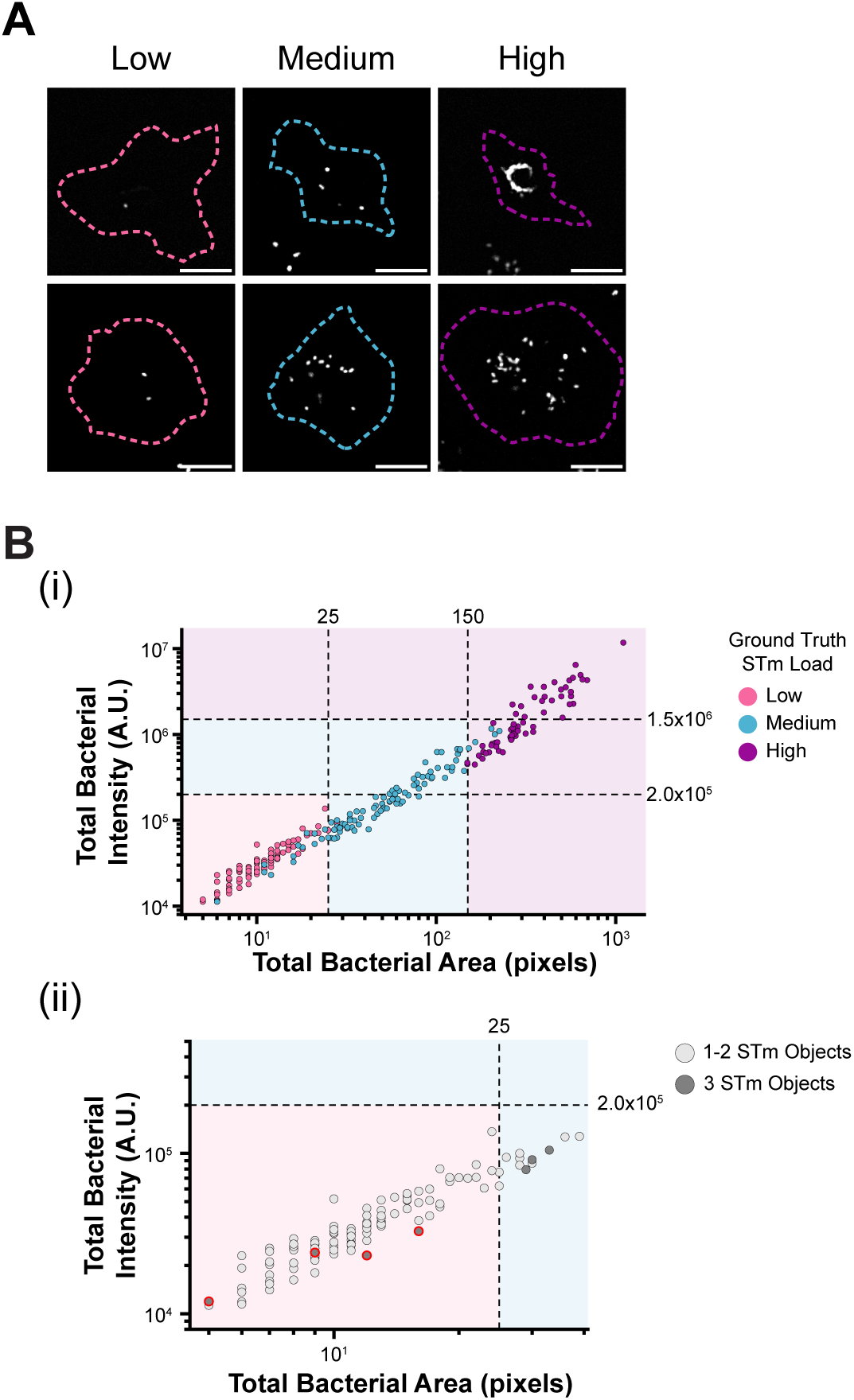
Classification of CIMs based on STm loads. (A) Representative images of infected CIMs with low (pink), medium (cyan), or high (purple) bacterial loads. Images show the CIM cytoplasm boundary (colored dashed line) and STm (grayscale). Scale bars=20 µm. (B) Classification strategy using features of bacterial objects. (i) Ground truth dataset generation and threshold definitions. A total of 254 STm-infected CIMs were manually classified as containing low (pink), medium (blue), or high (purple) bacterial loads. For each CIM, the total fluorescence intensity (arbitrary units, A.U.) and total area (pixels) of all intracellular bacterial objects combined was quantified. Using the manually classified dataset, total intensity and total area thresholds (dashed lines) were defined to classify CIMs into low (pink shaded region), medium (blue shaded region), or high (purple shaded region) STm load categories. (ii) Refinement using the number of bacterial objects. A subset of CIMs with medium STm loads in the ground truth dataset were initially misclassified in the low STm load category based on total intensity and total area thresholds. To rectify this, CIMs classified as low but containing >2 bacterial objects were reclassified into the medium load category. The plot shows total intensity and total area for a subset of CIMs from (i). CIMs with 1-2 bacterial objects are shown in light gray and CIMs with 3 bacterial objects are shown in dark gray. CIMs reclassified from low to medium based on the number of objects are highlighted with red outlines.

**Figure S4.2.**
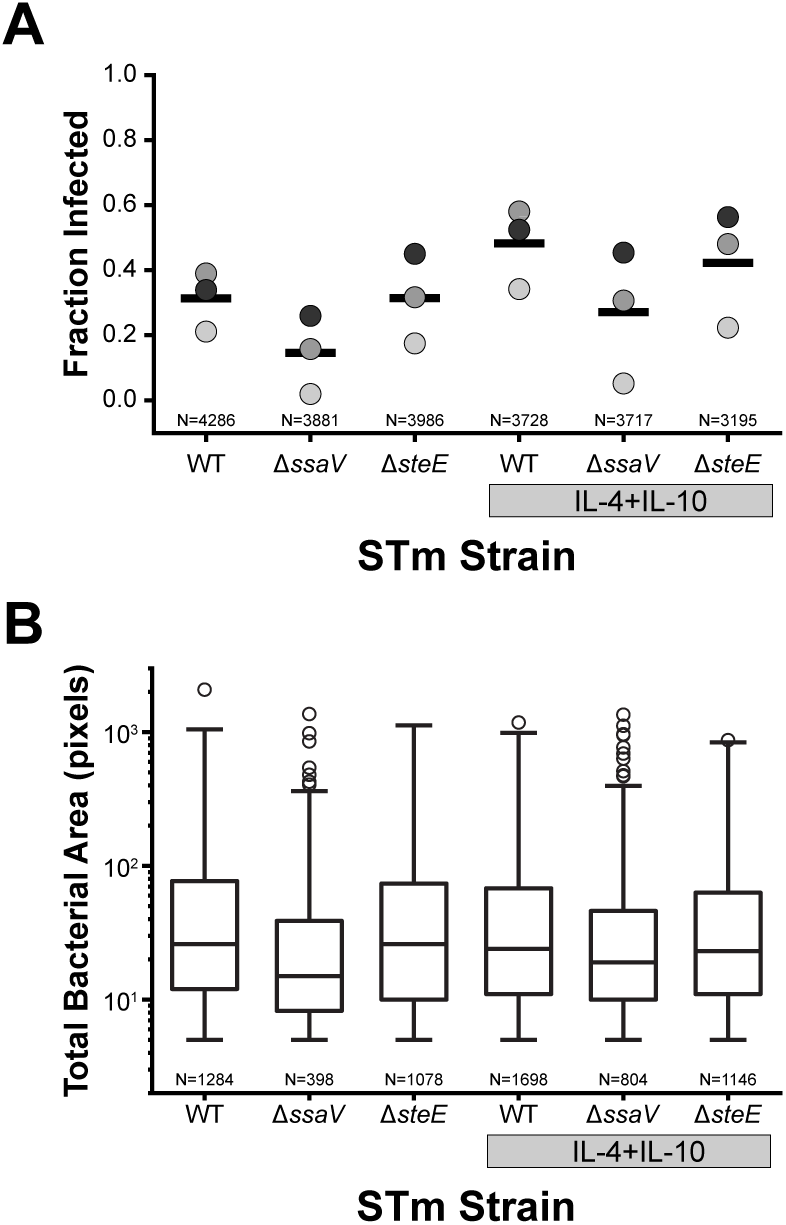
Infection frequency and bacterial replication of WT and mutant STm strains in CIMs. CIMs were exposed to WT, Δ*ssaV*, or Δ*steE* STm alone or in combination with M2 ligands (25 ng/mL IL-4 and 25 ng/mL IL-10) and fixed at 24 hours. Data correspond to Figure 4B. (A) Infection frequency. The fraction of infected CIMs (containing ≥1 bacterial objects) is shown for three biological replicates (shades of gray) with the mean shown as a black bar. (B) Bacterial replication. The total bacterial area (pixels) of infected CIMs was used as a proxy for intracellular bacterial replication. Data are pooled from three biological replicates. N=total number of CIMs. Results of statistical analyses are reported in Table S1. All pairwise comparisons vs WT STm had p>0.05 (not significant), Mann-Whitney U test with FDR correction.

**Figure S4.3.**
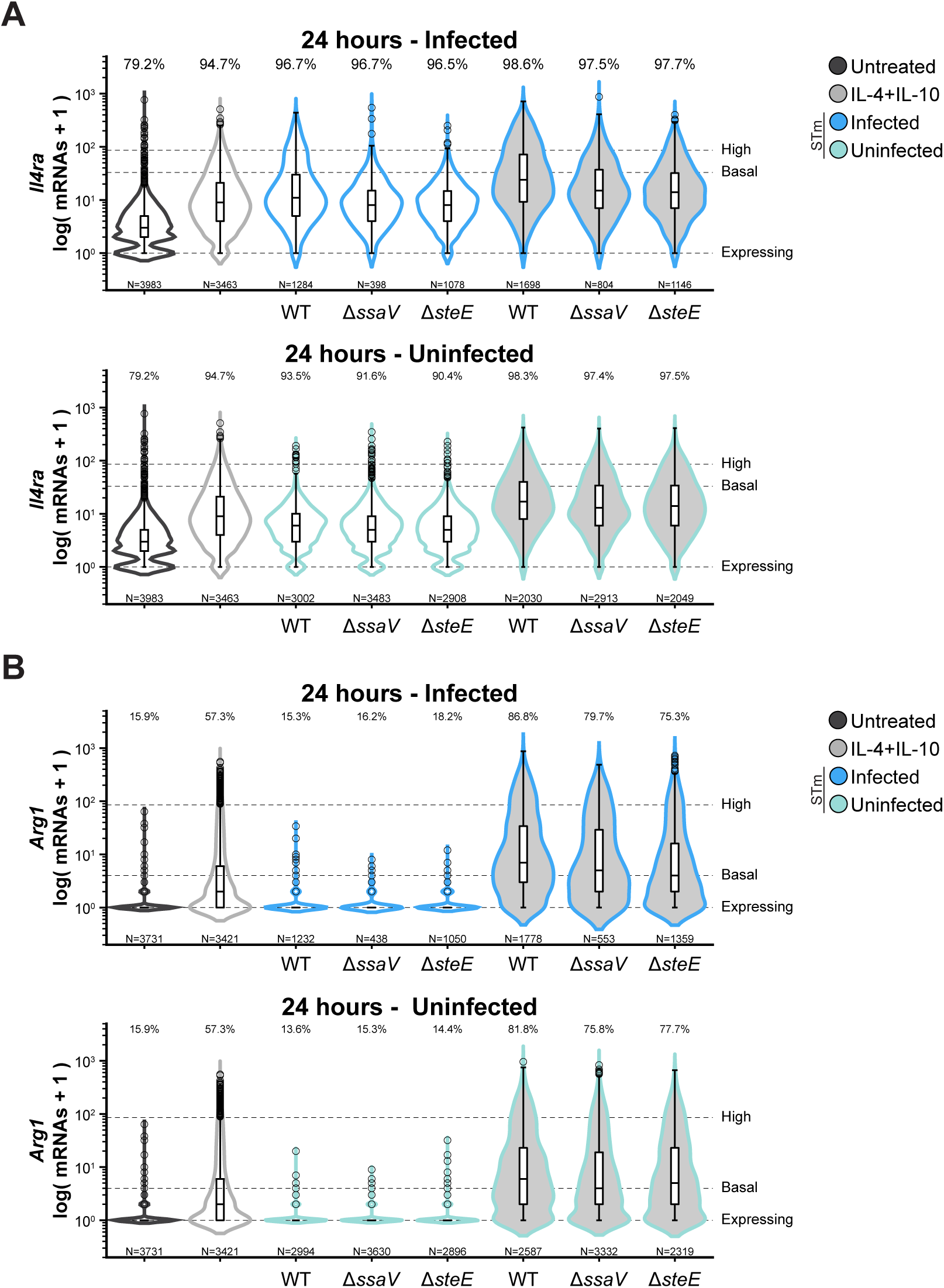
Il4ra and Arg1 gene expression change with STm effector secretion and the presence of M2 ligands. (A) *Il4ra* and (B) *Arg1* single-cell gene expression corresponding to data shown in Figure 4B. CIMs were left untreated, treated with M2 ligands (25 ng/mL IL-4 and 25 ng/mL IL-10), or exposed to STm (WT, Δ*ssaV*, or Δ*steE*) either alone (empty violins) or in combination with M2 ligands (filled violins). Samples were fixed at 24 hours, processed for smFISH, and imaged. CIMs were classified as infected (top) or uninfected (bottom) as described in Figure 1 (see Methods). For STm-exposed CIMs, the strain is indicated on the x-axis. Violin plots show transcript levels for *Il4ra* and *Arg1*, data are presented as log(mRNA +1), and the width of the violins represents the proportion of CIMs within the respective condition. Dashed lines indicate expressing CIMs (1 transcript), basal expression (32 mRNAs for *Il4ra* and 3 mRNAs for *Arg1*), and the 95^th^ percentile threshold (85 mRNAs for *Il4ra* and 84 mRNAs for *Arg1*) defining high-expressing CIMs. Percentages above the violins indicate the percentage of CIMs containing transcripts. Data are pooled from three biological replicates. N=total number of CIMs. Results of statistical analyses on smFISH data are reported in Table S1.

**Figure S4.4.**
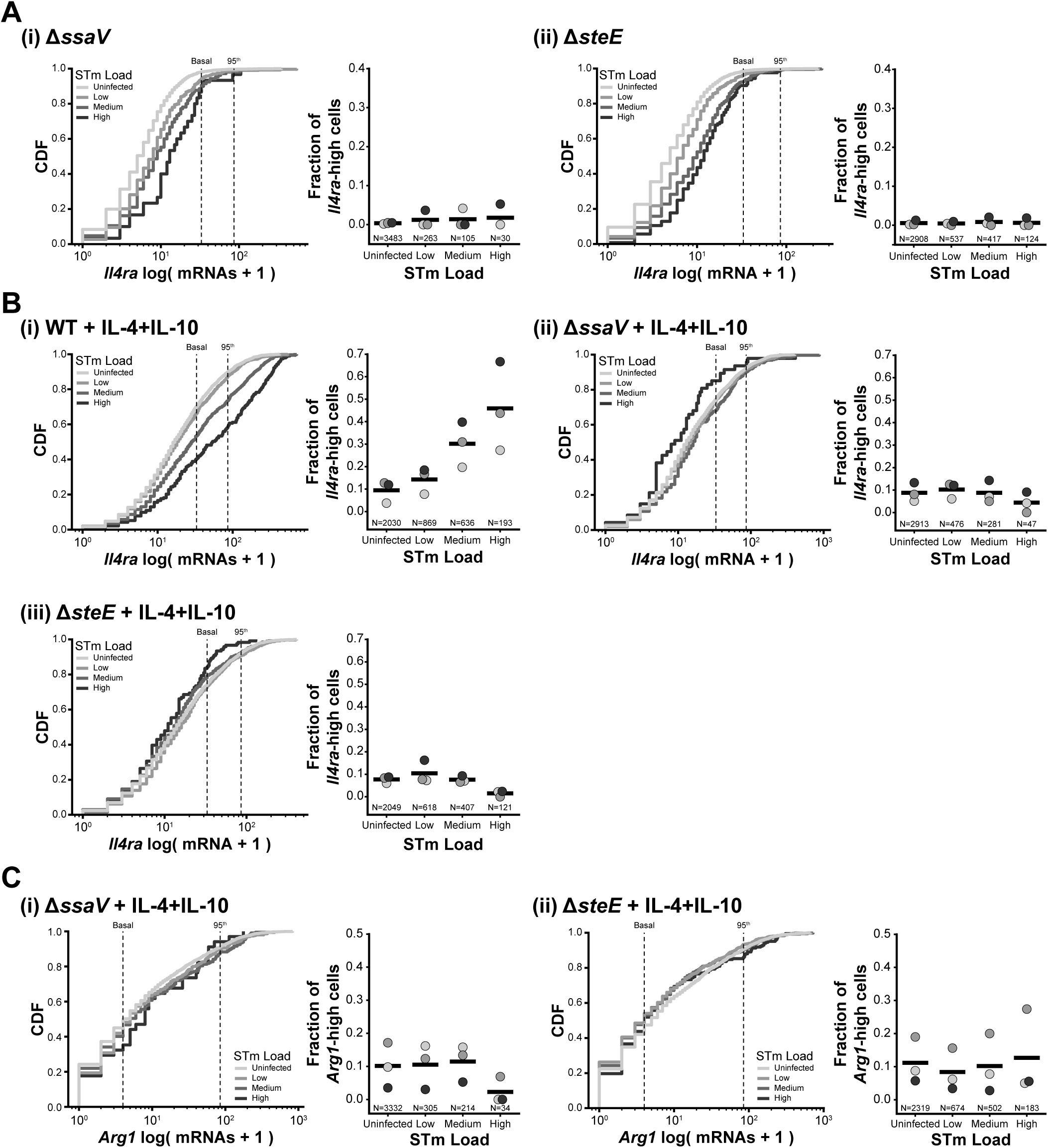
Effectors and M2 ligands affect the scaling of anti-inflammatory gene expression with STm load and the fraction of high-expressing CIMs. CIMs were exposed to STm strains (WT, Δ*ssaV*, or Δ*steE*) alone or in combination with M2 ligands (25 ng/mL IL-4 and 25 ng/mL IL-10). Data corresponds to that shown in Figure 4B. CIMs were fixed at 24 hours, processed for smFISH, imaged, and classified based on their STm load (see Figure S4.1 and Methods). (A) *Il4ra* expression in CIMs exposed to (i) Δ*ssaV* STm and (ii) Δ*steE* STm. (B) *Il4ra* expression in CIMs exposed to (i) WT STm and M2 ligands, (ii) Δ*ssaV* STm and M2 ligands, and (iii) Δ*steE* STm and M2 ligands. (C) *Arg1* expression in CIMs exposed to (i) Δ*ssaV* STm and M2 ligands and (ii) Δ*steE* STm and M2 ligands. For each subpanel: (Left) The cumulative distribution function (CDF) of mRNA counts plotted as log(mRNA +1) for each gene and pooled from three biological replicates. Different STm loads are shown in shades of gray. Dashed lines indicate basal expression (32 mRNAs for *Il4ra* and 3 mRNAs for *Arg1*) and the 95^th^ percentile threshold (85 mRNAs for *Il4ra* and 84 mRNAs for *Arg1*) defining high-expressing CIMs. (Right) The fraction of high-expressing CIMs (above the 95^th^ percentile) in each STm load category. Note that for comparative purposes, the scaling of the y-axes of (A) and (C) is consistent with the data shown in Figure 4A. The scaling of the y-axes of (B) was adjusted to account for the increase in the fraction of *Il4ra*-high cells in the WT STm and M2 ligands condition. Biological replicates are shown in shades of gray with the mean shown in black. N=total number of CIMs. Pairwise comparisons used a binomial generalized linear model (GLM) with post-hoc pairwise comparisons performed using linear contrasts on the fitted model coefficients, with Wald tests used to evaluate significance, with FDR correction. p>0.05 (not significant) for all pairwise comparisons except for (B) (i) *Il4ra*: WT STm and M2 ligands: all p<0.001 except uninfected vs low (p<0.05) and (C) (i) *Arg1*: (ii) Δ*steE* STm and M2 ligands: p<0.01 (low vs high). Results of statistical analyses on the fraction of *Il4ra*-high and *Arg1*-high across different bacterial loads are reported in Table S1.

**Table S1. Summary of data used for figures and statistical analyses.**

